# Profiling Transcription Initiation in Peripheral Leukocytes Reveals Severity-Associated Cis-Regulatory Elements in Critical COVID-19

**DOI:** 10.1101/2021.08.24.457187

**Authors:** Michael Tun Yin Lam, Sascha H. Duttke, Mazen F. Odish, Hiep D. Le, Emily A. Hansen, Celina T. Nguyen, Samantha Trescott, Roy Kim, Shaunak Deota, Max W. Chang, Arjun Patel, Mark Hepokoski, Mona Alotaibi, Mark Rolfsen, Katherine Perofsky, Anna S. Warden, Jennifer Foley, Sydney I Ramirez, Jennifer M. Dan, Robert K Abbott, Shane Crotty, Laura E Crotty Alexander, Atul Malhotra, Satchidananda Panda, Christopher W. Benner, Nicole G. Coufal

**Author notes:** Address correspondence to: Michael Tun Yin Lam, Division of Pulmonary, Critical Care, and Sleep Medicine, Department of Medicine, University of California San Diego, 9300 Campus Point Dr., #7381, La Jolla, California 92037, USA. authors contributed equally to this work (Sascha H. Duttke, Mazen F. Odish, Hiep D. Le, Emily A. Hansen, Celina T. Nguyen, Samantha Trescott, Roy Kim). both authors contributed equally to this work (Christopher W. Benner, Nicole G. Coufal).

## Abstract

The contribution of transcription factors (TFs) and gene regulatory programs in the immune response to COVID-19 and their relationship to disease outcome is not fully understood. Analysis of genome-wide changes in transcription at both promoter-proximal and distal cis-regulatory DNA elements, collectively termed the ’active cistrome,’ offers an unbiased assessment of TF activity identifying key pathways regulated in homeostasis or disease. Here, we profiled the active cistrome from peripheral leukocytes of critically ill COVID-19 patients to identify major regulatory programs and their dynamics during SARS-CoV-2 associated acute respiratory distress syndrome (ARDS). We identified TF motifs that track the severity of COVID- 19 lung injury, disease resolution, and outcome. We used unbiased clustering to reveal distinct cistrome subsets delineating the regulation of pathways, cell types, and the combinatorial activity of TFs. We found critical roles for regulatory networks driven by stimulus and lineage determining TFs, showing that STAT and E2F/MYB regulatory programs targeting myeloid cells are activated in patients with poor disease outcomes and associated with single nucleotide genetic variants implicated in COVID-19 susceptibility. Integration with single-cell RNA-seq found that STAT and E2F/MYB activation converged in specific neutrophils subset found in patients with severe disease. Collectively we demonstrate that cistrome analysis facilitates insight into disease mechanisms and provides an unbiased approach to evaluate global changes in transcription factor activity and stratify patient disease severity.

## Introduction

Acute respiratory distress syndrome (ARDS) is the cardinal clinical feature and the primary contributor to mortality in severe SARS-CoV-2 infection. Cytokine and single-cell analyses support a model for a prolonged hyperinflammatory response driving diffuse alveolar damage (1–8). Tremendous effort has been invested in repurposing available therapeutics for the treatment of severe COVID-19. To this end, glucocorticoids and anti-interleukin-6 receptor monoclonal antibodies reduce mortality for severe cases of COVID-19. Importantly, the mortality benefits were only observed in subgroups of patients. Therapies provided to inappropriate patient subpopulations may cause harm (9). These results signal the unmet need for additional therapeutic targets and novel stratification strategies to identify the right patient for the right therapy.

Understanding the dynamic relationship of transcription factor activity and disease severity may offer insights for precision therapy. Transcriptional responses are an important component of the host response to infectious disease. Not only do SARS-CoV-2 infected cells up-regulate antiviral gene expression programs to halt the viral spread, but they also signal to activate regulatory networks in other cells and tissues to mount a coordinated immune response to the pathogen. Transcription factors (TFs) are vital in orchestrating these transcriptional responses. TFs integrate upstream inflammatory and immunological signals to direct changes in the transcriptional programs mediating cellular adaptation and function. TFs bind DNA *in cis* at regulatory regions through specific recognition DNA sequences, also known as TF motifs.

Unbiased, genome-wide profiling of cis-regulatory elements, coupled with computational analysis for motif enrichment, is a powerful tool to discover transcriptional regulatory mechanisms. Understanding what these pathways and cell types are, how they vary across individuals and time, and how they are dysregulated in severe disease is critical to understanding COVID-19 and host response to sepsis and severe lung injury.

This study used unbiased cistrome profiling of peripheral leukocytes from COVID-19 patients to decipher regulatory networks activated or repressed during severe SARS-CoV-2 infection. Given that transcription is a hallmark of regulatory activity from promoters and enhancers (10), we used capped small (cs)RNA-seq to measure the transcriptional activity from genome-wide cis-regulatory elements. csRNA-seq captures short, 5’ capped RNAs (20-60nt) associated with engaged RNA polymerase II and defines the transcription start sites at a single-nucleotide resolution of both stable and unstable transcripts such as enhancer RNA (eRNA) (11). Moreover, csRNA-seq focuses motif analysis on the active cistrome compared to the assessment of chromatin accessibility, where regulatory elements may be accessible but transcriptionally inactive (e.g., open-poised, insulators, etc.) (12).

We performed csRNA-seq with matched samples for RNA-seq to capture the steady-state transcriptome and cytokine analysis on peripheral leukocytes to construct a natural history of TF programs in ARDS associated with severe COVID-19. We profiled the active cistrome from 22 individuals, with a median of 7 time points encompassing early, mid, and late hospitalization. Our analysis revealed regulatory programs associated with specific TFs and cell types that exhibited activity patterns correlated with clinical phenotypes. We identified a role for inflammatory transcription factor families in early-stage disease, including Nuclear Factor-kappa B (NFkB), Signal Transducer and Activator of Transcription (STAT), and Interferon Regulatory Factors (IRF). We also identified robust disease associations for other TFs and TF families, including Glucocorticoid Receptor, Nuclear Factor E2 Related Factor 2 (NRF2), E2F, MYB, and the family of microphthalmia (MiT/TFE). Because our dataset provides precise genomic locations of regulatory activity during COVID-19 infection, we cross-referenced the active cistrome with existing genomics data, including chromatin state maps and genetic variants associated with COVID-19 clinical outcome. We identified significant enrichment of diseased-associated single nucleotide polymorphisms (SNPs) in distinct TF regulatory networks. Using target gene expression as a measure of TF regulatory network activity, we independently stratified COVID-19 patients with poor outcomes in a large published cohort with early admission transcriptomics. Patients with high expression for E2F/MYB and STAT targets, or target expression that is low for Type 1 interferon and high for STAT targets, had the most severe outcome. Integrating these findings with published single-cell RNA-seq, we show that dysregulated E2F/MYB, STAT, and Type 1 interferon signals are reflected in the differential distribution of neutrophil subsets. These findings showcase the utility of profiling transcription initiation to reveal regulatory programs from blood samples and provide insight into the key TFs and pathways activated during the host response to COVID-19.

## Results

### Transcription initiation analysis with csRNA-seq reveals the active cistrome from peripheral leukocytes during severe SARS-CoV2 infection

Genomic regions that initiate transcription represent active cis-regulatory elements (10, 13, 14). To understand the dynamic changes in cistrome activity during the course of COVID- 19, we profiled transcription initiation events from peripheral leukocytes of COVID-19 patients (Fig 1a). We isolated peripheral blood from 5 healthy controls and 17 patients, with a median of 7 time points per patient, spanning a median hospitalization of 8 days (range 1-22 days), for a total of 92 time points. 16 of the 17 patients required care in the Intensive Care Unit (ICU). 13/17 required mechanical ventilation. 48% of the samples were collected from patients with severe lung injury, defined as a Modified Murray Score > 2.5 (15). Five patients recovered and were discharged within ten days (Fast Recovery). Nine survivors had prolonged hospitalization with three in-hospital fatalities (Fig 1b, Supp Table 1).

**Figure 1.**
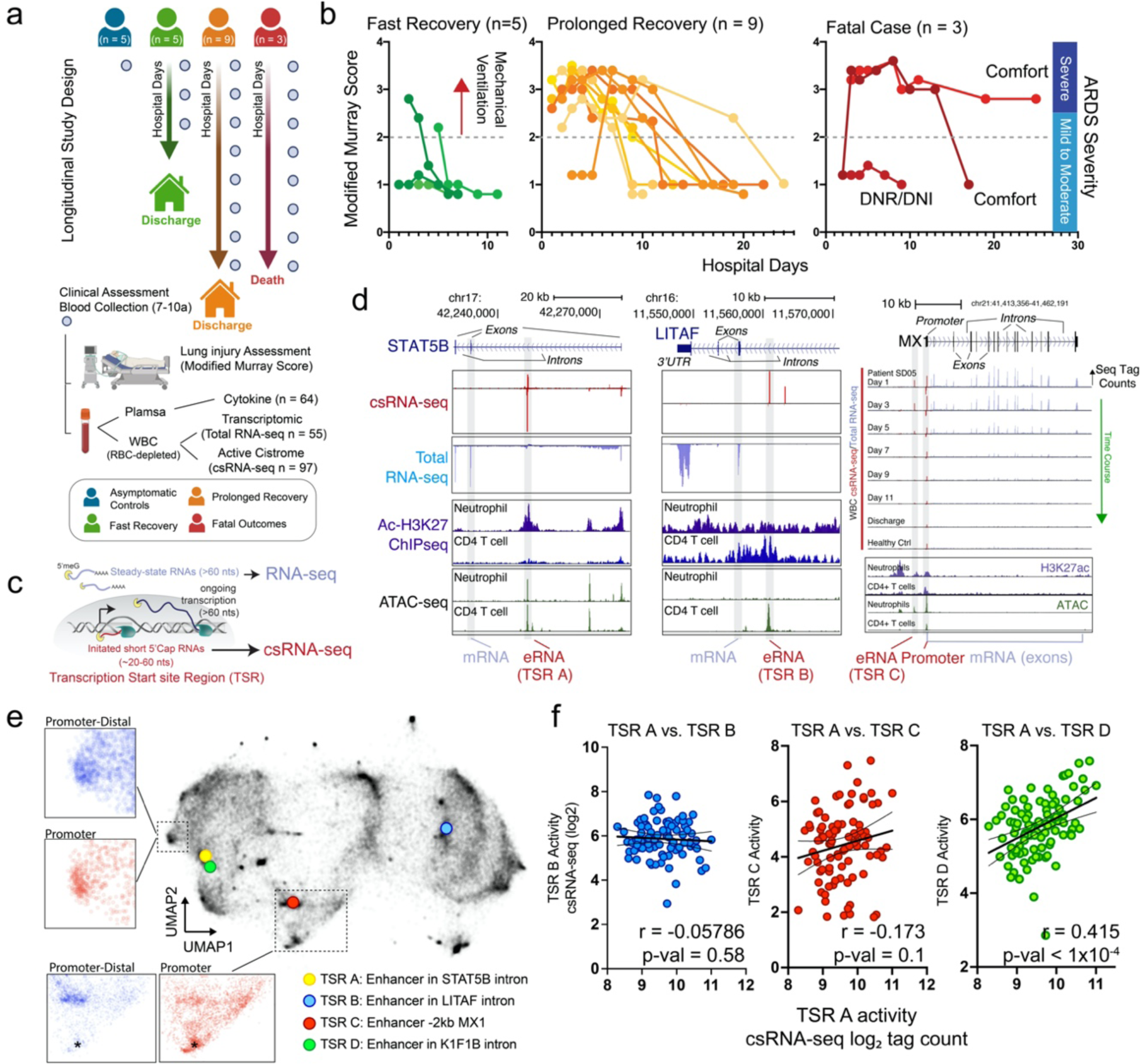
Activated immune cistrome from peripheral leukocytes of hospitalized COVID-19 patients. a-b. Longitudinal study design sampled plasma and peripheral leukocytes of hospitalized COVID-19 patients across different stages of lung injury quantified by the Modified Murray Lung Injury Score. Ninety-seven samples were included for active cistrome analysis using capped-short RNA-seq (csRNA-seq). For fatal cases, the collection ended after patients transitioned to comfort care. One patient declined resuscitation or intubation (DNR/DNI). c) csRNA-seq captures short 5’ capped RNA species, including active cis-regulatory elements and gene promoters, collectively termed Transcription Start site Regions (TSRs). d) csRNA-seq identifies transcriptional activity (red) in putative enhancers (eRNAs) in the *STAT5B* (left), *LITAF* (middle) and MX1 (right) loci. The *STAT5B* enhancer resides in a neutrophil-specific active chromatin region (+ acetylated H3K27 and + ATAC-seq), whereas the *LITAF* enhancer resides in CD4 T cell active chromatin region. *MX1* eRNA correlates with *MX1* gene expression over time. e) Unbiased clustering of 93,465 TSRs grouped by similarity of their activity across 97 samples using Uniform Manifold Approximation Projection (UMAP). Inset shows TSRs residing in the promoter (red) or promoter-distal regions (blue). f) Pearson correlation of csRNA-seq levels from TSR A compared to TSR B (left), TSR C (middle), and TSR D (right).

Across a total of 97 csRNA-seq libraries, we cumulatively identified 93,465 genomic regions with ample evidence of transcription initiation, termed Transcription Start site Regions (TSRs). >95% of the identified TSRs overlapped with open chromatin regions defined by ATAC-seq in one or more leukocyte cell populations (Supp Fig 1) (16, 17). 42% of the TSRs mapped to the vicinity (< 500 bp) of annotated gene promoters. Nucleotide frequency analysis relative to transcription start sites showed defined features consistent with promoter elements. 58% of TSRs mapped to promoter-distal regions co-localized with epigenetic marks in leukocytes associated with active enhancers (18), with nucleotide frequency distinct from promoters (Supp Fig 1a-c). For example, the intronic regions of the *STAT5B* and *LITAF* loci have open chromatin regions with transcription initiation activity consistent with enhancer elements (Fig 1d left, middle panel). The interferon-induced *MX-1* gene has two TSRs representing the gene promoter and enhancer (Fig 1d right panel). The *MX-1* enhancer TSR resides 2kb upstream of the promoter in an open chromatin region (ATAC-seq) surrounded by modified histones (acetylated-H3K27) consistent with an active neutrophil-specific enhancer. Notably, the activity of the enhancer TSR correlates with the transcriptional initiation signal and the stable RNA transcript level (total-RNA seq) of *MX-1*. Overall, we generated a dataset of cistromic activity from peripheral leukocytes that accurately identify promoters and enhancers in leukocytes from patients with severe SARS-CoV-2 infection.

### Agnostic TSR clustering contextualizes the activity of the immune cistrome with lung injury severity

The activity of the cistrome is influenced by multiple biological factors, including regulation by TFs, cell-type specificity, disease severity, and other physiological factors. To appreciate these interactions, we performed unsupervised clustering based on TSR activity across all 97 samples capturing TSRs co-regulated across patients at different time points and disease severity states. TSRs regulated by a common biological mechanism(s) should display similar activity and therefore cluster together. To this end, we applied hierarchical clustering, revealing 26 distinct TSR clusters, and used Uniform Manifold Approximation and Projection (UMAP) to group TSRs by their similarity and visualize them in 2D space (Fig 1e, Suppl Fig 2).

**Figure 2.**
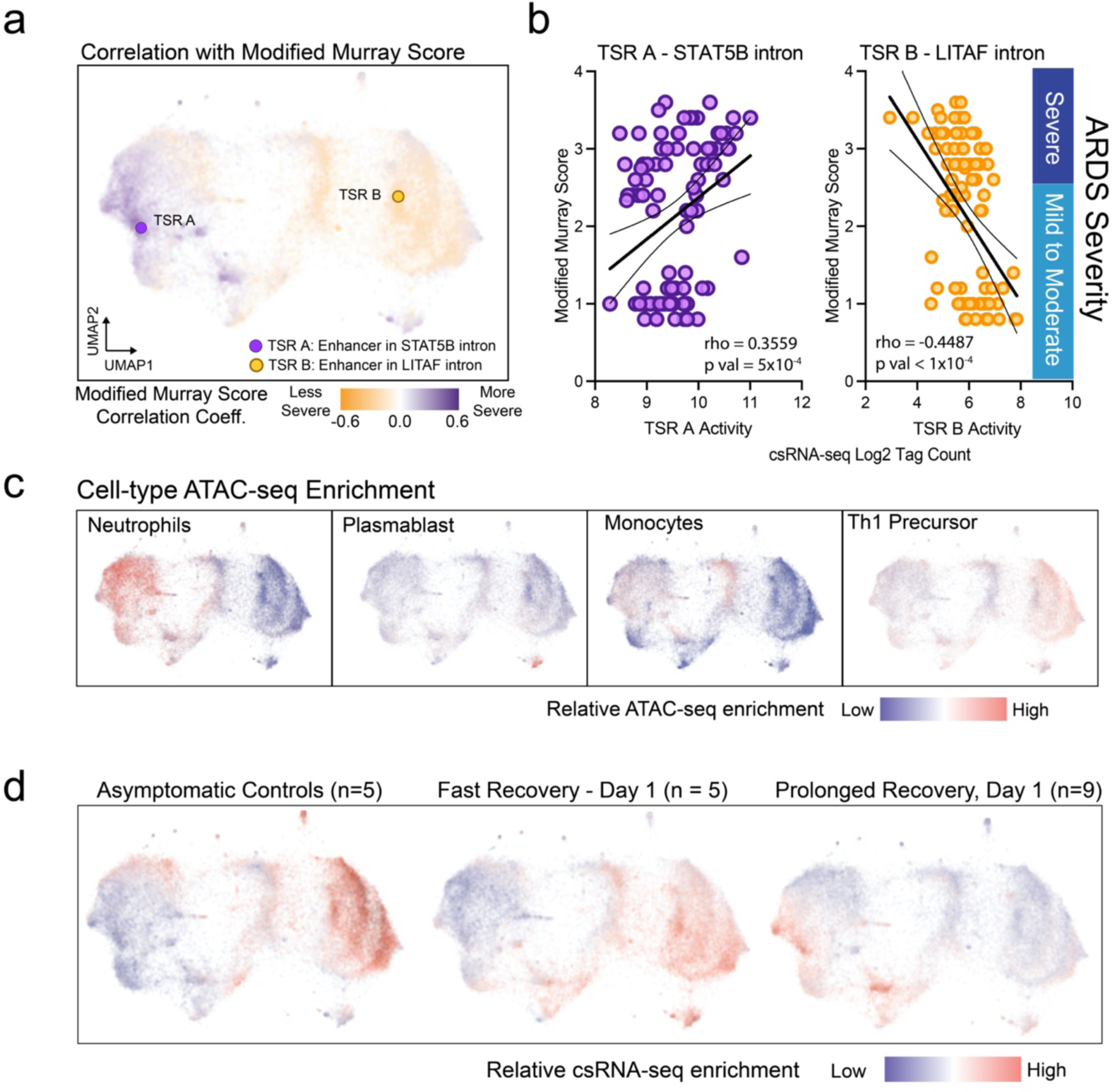
Differential activation of the immune cistrome at different disease states. A) UMAP of TSRs, shaded based on the correlation of their activity profile with lung injury score B) Spearmen correlation analysis of the activities of TSR A (left) and TSR B (right) with modified Murray Lung Injury Score. C) Open-chromatin ATAC-seq enrichment in neutrophils, plasmablasts, monocytes, and Th1 precursor lymphocytes from each TSR visualized on the immune cistrome UMAP (16, 17). Red delineates high ATAC-seq enrichment relative to other hematopoietic cell types. D) Genome-wide relative average TSR activity in asymptomatic controls (n=5) and patients with fast (n=5) or prolonged recovery (n=9) on the first day of enrollment.

We first interrogated the relationship of cistrome activity with disease severity. We focused on lung injury and quantified how individual TSR activity correlated with patients’ Modified Murray Lung Injury Score. By coloring the TSR UMAP by their correlation to lung injury, we observed that TSRs were primarily segregated by disease severity (Fig 2a). For example, TSR A found in a *STAT5B* intronic region has an activity that correlates with lung injury (rho = 0.3559, p-value = 5x10^-4^). In contrast, TSR B located in an intron of *LITAF* has a negative correlation to lung injury (rho = -0.4487, p-value < 1 x 10^-4^) (Fig 2b). Because each sample profiles the entire bulk population of peripheral leukocytes, we next estimated the cell type specificity of each TSR to compare with the lung injury spectrum. We utilized a hematopoietic cell-type-specific reference ATAC-seq dataset from healthy donors to identify discrete cistrome clusters associated with neutrophils, monocytes, lymphocytes, and plasmablasts specific peaks (Fig. 2c, Supp Fig 3) (16, 17). Lymphocyte-associated TSR- clusters correlated with lower Modified Murray Lung Injury Scores, consistent with the observation that the neutrophils to lymphocytes ratio is elevated in severe COVID-19 (19). Of significance, not all TSRs within the neutrophil clusters are positively correlated with lung injury, suggesting different transcriptional programs may be activated in neutrophils associated with different disease states. Furthermore, TSRs with high activity on the first day of enrollment are distinct between patient severity groups, collectively suggesting the active cistrome encodes valuable information about disease states (Fig 2d).

**Figure 3.**
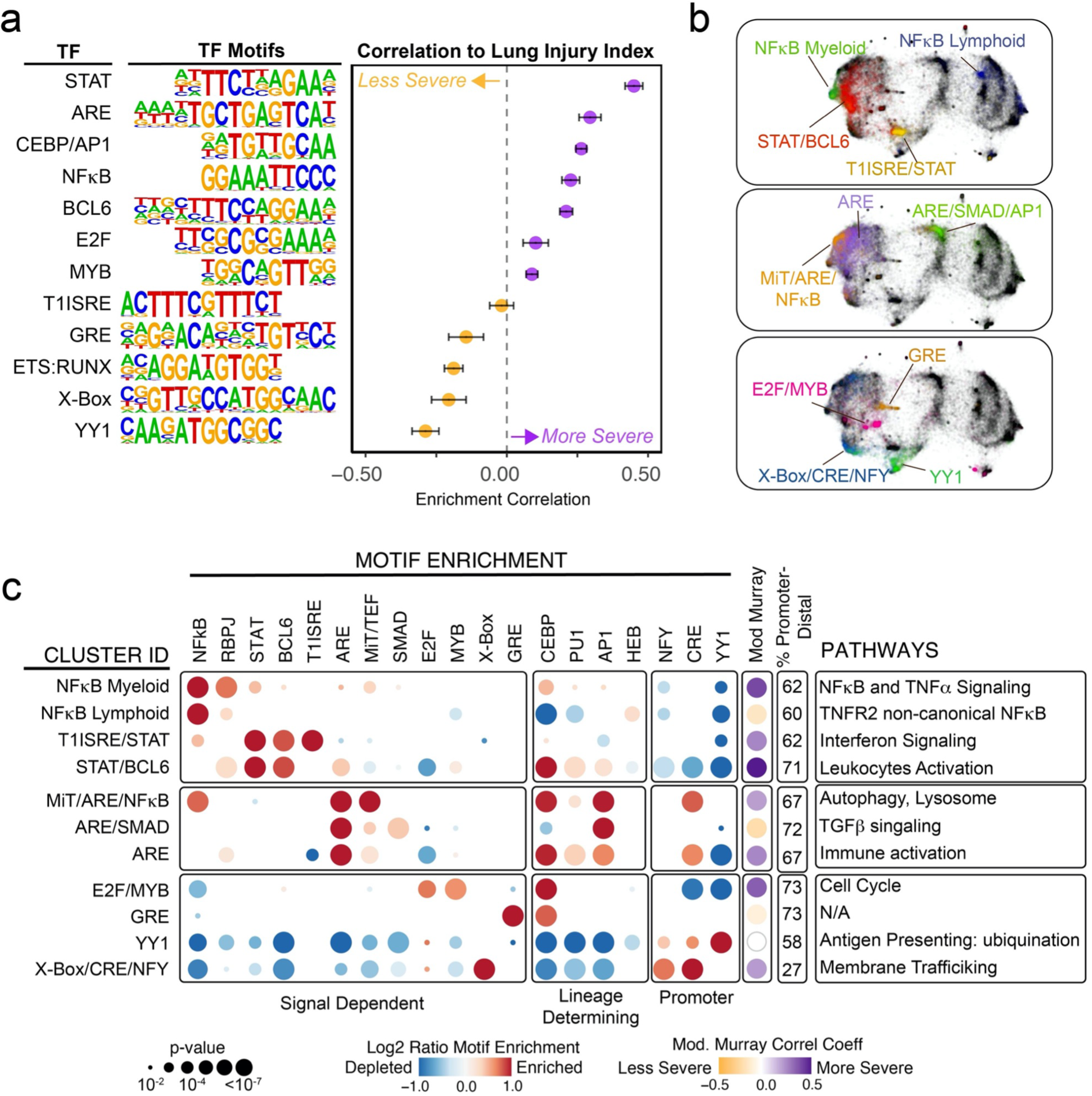
Distinct cistrome clusters identify co-enrichment of transcription factor (TF) motifs suggestive of co-regulatory mechanisms. a) A logistic regression analysis (MEIRLOP) identified transcription factor (TF) motifs enriched in regulatory elements associated with high (violet) or low (orange) lung injury indices. Each dot represents the enrichment coefficient of TF motifs in TSRs with activity profiles highly correlated with the lung injury index. Error bars represent the lower and upper 95% confidence intervals. The enrichments of all motifs, except for T1ISRE, are all statistically significant (adj. p < 0.0001). b) UMAP representation showing discrete TSR clusters labeled by representative TFs exhibiting the highest enrichment in each cluster. c) Motif analysis depicts co-enrichment of signal-dependent, lineage-determining, and promoter TF motifs. Red depicts the Log2 ratio enrichment of the motif frequency in the TSR cluster relative to all TSRs; blue, depletion. The dot size represents the Fisher Exact Test p-value. Functional enrichment/GO analysis identifies top pathways from genes associated with each TSR cluster.

We then investigated the association of transcription factor motifs in the active cistrome in relationship to disease severity. We searched for known TF motifs within -150 to +50 bp of the transcription start sites and performed a score-based logistic regression model (MEIRLOP) to identify TF motifs associated with lung injury (Fig 3a) (20). We identified enrichment of STAT, NFkB, and myeloid lineage determining (LD) CEBP/AP1 TF motifs in TSRs associated with severe lung injury. Interestingly, the Antioxidant Responsive Element (ARE) is one of the top five motifs most associated with severe lung injury. This motif is recognized by transcription factors involved in reduction-oxidation homeostasis, including the members of NRF, the small- MAF, and the BTB and CNC Homology (BACH) families. Genetic variants in the NRF member *NFE2L2* are associated with higher susceptibility to ARDS (21), thus providing biological plausibility that this pathway is active in COVID-19 ARDS. TSRs associated with low lung injury indices exhibited enrichment in the YY1 promoter element, glucocorticoid response element (GRE), X-box, and motifs recognized by lymphocytes lineage determining TFs (LDTFs) including ETS/RUNX (Fig. 3a). The correlation of GRE motifs with lower lung injury index is consistent with studies showing the benefit of glucocorticoid therapy in severe COVID-19.

### Identification of a cooperative transcriptional factor regulatory network

To further define transcriptional regulatory mechanisms underlying cistromic activity, we performed motif enrichment and pathway analysis on TSRs that segregated into distinct clusters, representing co-regulated networks of TSRs (Fig 3b-c, Supp Fig 4). This analysis successfully captured enrichment of key immune regulators in specific clusters, suggesting we can segregate and track the activity of distinct pathways across our dataset. Furthermore, motif analysis identified co-enrichment of signal-dependent TFs (SDTFs) with LDTFs, consistent with the model that pioneering TFs establish accessible sites for cell-type-specific transcriptional regulation (22). This point is exemplified by the two distinct clusters with enrichment for NFkB motifs (Fig 3b-c, top panel, Supp Fig 4). One NFkB cluster is myeloid-centric with 904 TSRs, co-enriched for CEBP, and strongly correlated with severe lung injury. 62% of the TSRs are in enhancers. TSRs located in promoters of protein-coding genes within this cluster are associated with the canonical pathway of NFkB and TNFα signaling in gene ontology analysis. The second NFkB cluster is lymphocyte-centric, negatively correlated with lung injury, with 1,136 TSRs and motif co-enrichment for the E-protein HEB, critical for T cell development (23).

**Figure 4.**
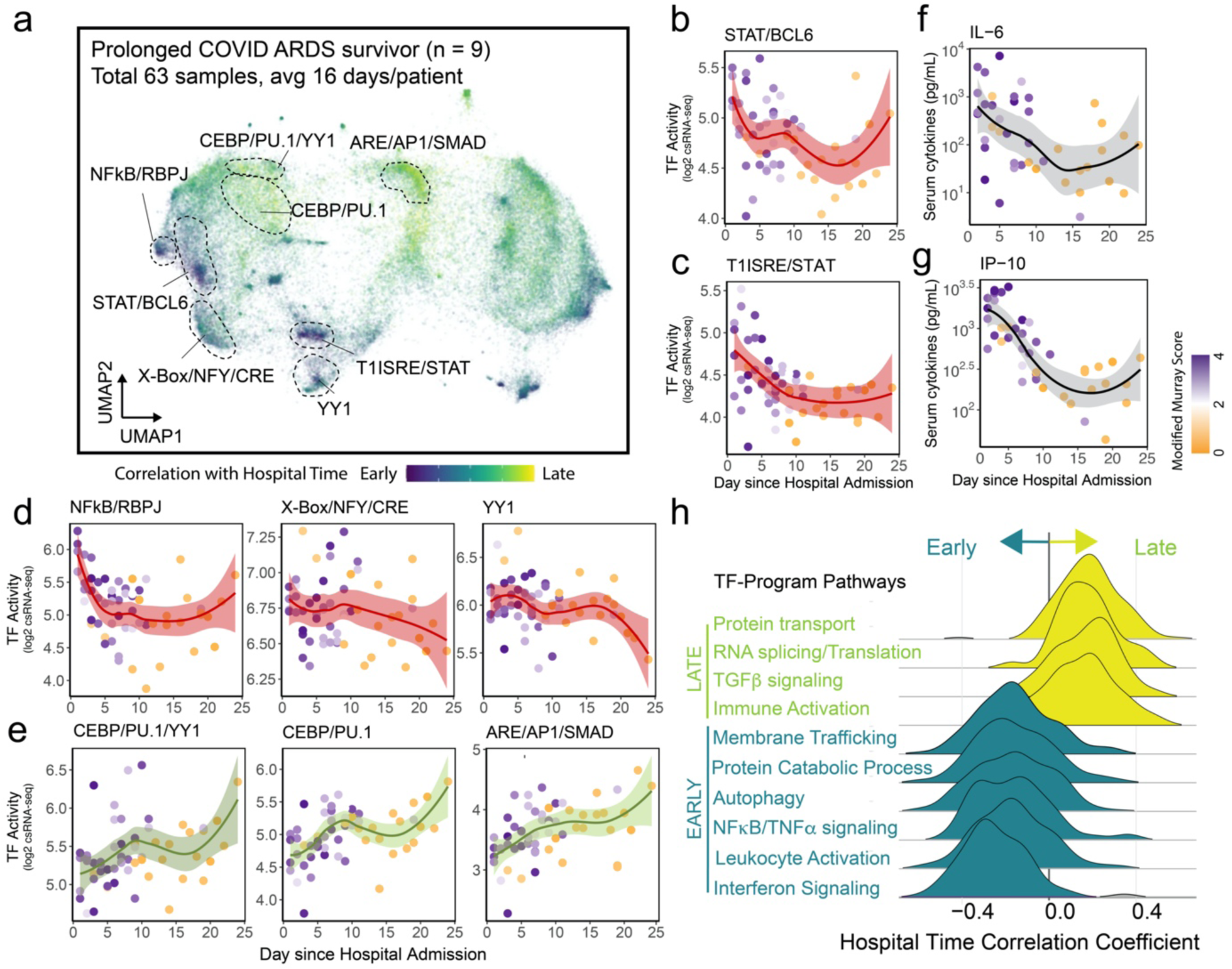
The natural progression of transcriptional programs during the clinical course of COVID19 ARDS. a) Genome-wide kinetic correlation analysis for cistrome activity to hospital days in critically ill COVID19 ARDS survivors (n = 9) with prolonged recovery (the median number of time points is 7 per patient; total = 63 samples). Correlation coefficients for each TSR activity to hospital time are overlaid on the UMAP. Purple indicates higher activity early in the hospital course; yellow, later hospital course. b-e) Time course of TF-activity in clusters enriched for (b) STAT/BCL6, (c) T1ISRE/STAT, (d) NFkB/RBPJ, X-Box/NFY/CRE, and YYI, e) ETS/YY1, CEBP/PU1, and ARE/AP1/SMAD. TF activity represents the median log2 csRNA-seq signals of all TSRs in a given cluster. f-g) Serum cytokines for (f) IL6 and (g) IP-10 implicated in the STAT and Interferon pathway, respectively. Each point represents the median log2 of TSR cluster activity with the color indicating the lung injury score at those time points (violet = high; gold = low). The line and shaded region correspond to the smooth conditional mean and 95% confidence intervals, respectively. h) Gene pathways enriched in the early (purple) and late (yellow) TF programs. Ridge plot shows the time-TSR activity correlation coefficient of genes in the respective pathways.

To test the notion that our analysis distinguishes NFkB regulatory programs activated in different cell types, we cross-examined NFkB p65/RELA chromatin localization in activated myeloid (24) and lymphoid cells (25) from publicly available Chromatin Immunoprecipitation (ChIP)-seq studies (Supp Fig 5). The TSRs from the myeloid NFkB cluster have a more significant overlap with NFkB ChIP-seq signal from activated monocyte-derived macrophages than from activated CD4+ T cells. In contrast, the lymphoid NFkB cluster has a greater overlap with NFkB ChIP-seq signal from activated CD4+ T cells (Supp Fig 5b), demonstrating that motif analysis of co-regulated TSRs can provide insights about activated pathways, their TFs, and cell types of activity.

**Figure 5.**
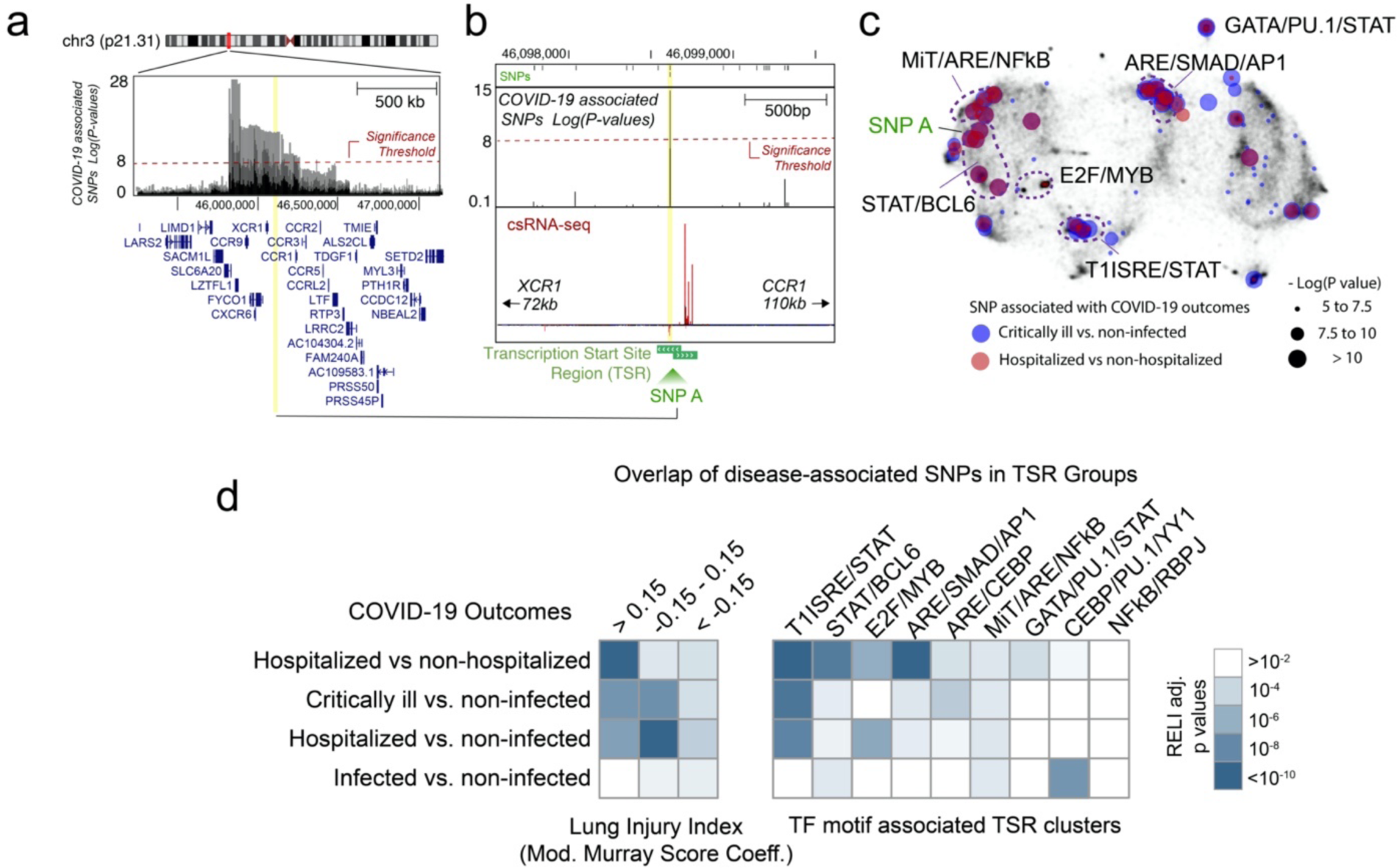
Distinct TSR clusters exhibit significant enrichment of single nucleotide polymorphisms associated with COVID-19 clinical outcome. a) The *LZTFL1* locus in chromosome 3p21.31 harbors numerous SNPs associated with COVID-19 clinical outcome (p-value < 5 x 10^-8^). SNP A (rs34460587, -log p-value > 15, hospitalization vs. non-hospitalized COVID-19 cases) lies within -300 to + 100 bp of a transcription start sites (TSS) located in the intergenic region between the *CCR1* and *XCR1* genes. c) UMAP showing the distribution of TSRs with COVID-19 associated SNPs overlap. d) Statistical analysis for enrichment of COVID-19 associated SNPs in TSRs based on lung injury index (left) or TSR clustering (right) using Regulatory Element Locus Intersection (RELI) (31). The color represents the RELI corrected p-values that account for the underlying genetic structure.

Interestingly, we often observed co-enrichment of multiple SDTFs within the same TSR cluster, suggesting coactivation by multiple regulators or pathways. This is exemplified by a cluster with a Type-1 interferon sensitive responsive element (T1ISRE) signature, which exhibited co-enrichment for the STAT motif, consistent with the role of STAT in interferon signaling during viral infection (Fig 2b-c top panel) (26).

The redox-responsive ARE motif participated in several distinct TSR clusters co-enriched with multiple SDTF motifs representing unrecognized SDTF-SDTF regulatory networks (Fig 3b-c, middle panel). First, an enhancer-centric 1,296-TSRs neutrophil cluster (Supp Fig 3-4) exhibited co-enrichment for motifs recognized by the MiT/TFE family (MITF, TFE3, TFEB, and TFEC) and NFkB. Genes located in the vicinity of these TSRs were functionally enriched for autophagy, lysosomes, and membrane trafficking, consistent with the role of the MiT/TFE family in reprogramming metabolism during stress (Fig 3c, middle panel) (27). In this cluster, TSRs containing both ARE and MiT motifs are enriched 3.4-fold relative to other active TSRs (Chi-square p-value < 1.0 x 10^-5^, two-tailed), suggesting a model where MiT/TFE and the NRF/small- MAF/BACH family are acting on the same regulatory regions (Supp Fig 6).

**Figure 6.**
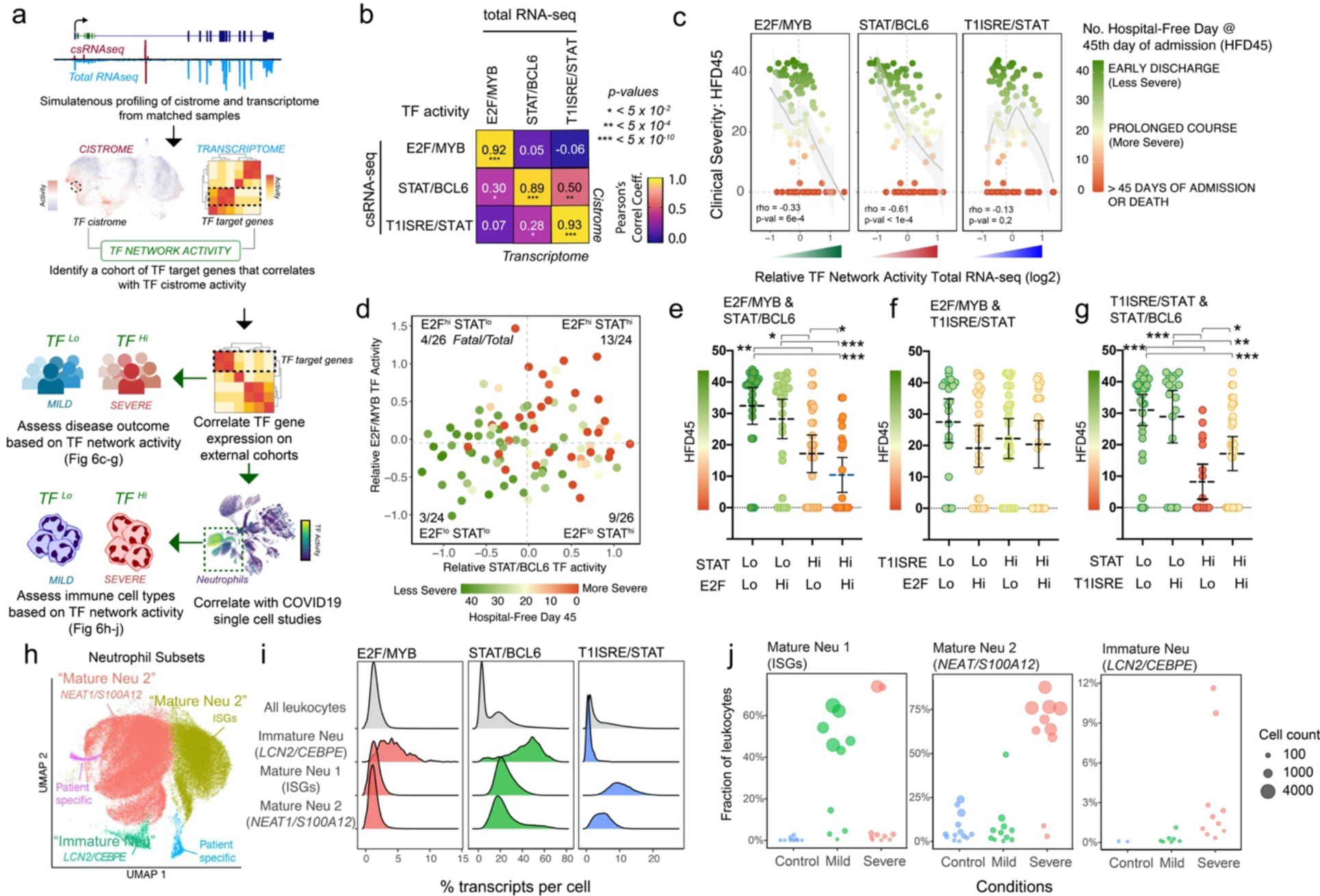
Dysregulated E2F/MYB, STAT/BCL6, and T1ISRE/STAT programs are associated with severe COVID-19. a) Framework for applying aggregate TF-network target gene expression as a representation of TF activity to validate cistrome-disease relationships in patient clinical outcome and cellular subtypes. b. Pair-wise Pearson’s correlation analysis of TF activity as determined by cistrome (csRNA-seq) and target gene expression (total RNA-seq) for E2F/MYB, STAT/BCL6, and T1ISRE/STAT using 55 matched samples. Correlation coefficients and the p-values for each pair-wise comparison are indicated. c. Spearman correlation of clinical severity (HFD45) with E2F/MYB, STAT/BCL6, and T1ISRE/STAT activity in an external COVID19 cohort (n = 100) (32). Smooth conditional means and 95% confidence intervals are depicted. d) Scatterplot of individual patient samples from the external COVID19 cohort based on STAT/BCL6 (x-axis) and E2F/MYB (y-axis) activity. The color of each point represents clinical disease severity (Red = severe; green = mild). The dash lines demarcate the medians for STAT/BCL6 (vertical) and E2F/MYB (horizontal) activity. The fatal cases for each quadrant are significantly different (Chi-square, p-value = 0.004, two-tailed). e-g) The average HFD45 for patient subgroups based on the activity of (e) E2F/MYB and STAT/BCL6, (f) E2F/MYB and T1ISRE/STAT, and (g) T1ISRE/STAT and STAT/BCL6. Error bars represent 95% confidence intervals from the mean (horizontal lines). One-way ANOVA with multiple comparisons shows statistical differences between subgroups. Adjusted p-values * < 0.05, ** < 0.005, *** < 0.0005. h) Three neutrophil subsets identified from COVID19 peripheral leukocyte single-cell RNA-seq analysis (8). i) Distribution of activities in E2F/MYB, STAT/BCL6, and T1ISRE/STAT programs per cell in each neutrophil subset. j) The cellular distribution of neutrophil subsets in control and COVID19 patients with mild and severe disease.

A second ARE cluster is enhancer-centric with 1,235 TSRs overlapping monocyte open chromatin (Fig 3b-c, middle panel; Supp Fig 3-4). In addition to ARE, this cluster had motif enrichment for Activator Protein 1 (AP1) and SMAD, the transducers for the Transforming Growth Factor β (TGFβ) signaling pathways. In addition, the target genes within this cluster were associated with TGFβ signaling in gene ontology analysis (Fig 3c, middle panel).

Consistent with this assertion, genome-wide TF localization studies in macrophages found that Nfe2l2, Smad3, and AP1 member Fos co-localized in common regulatory regions upon exposure to tissue damage signals (28).

We identified co-enrichment for the cell cycle and proliferation E2 Factor (E2F) and MYB TF families in a cluster of 2,364 TSRs (Fig 3b-c, bottom panel). The activity of this cluster was highly associated with lung injury severity (correlation = 0.36). When assessed individually using rank-based logistic regression, E2F and MYB motifs had a weak association to lung injury indices (Fig 3a, MEIRLOP coefficients = 0.102 and 0.077, respectively). Indeed, the E2F motif was enriched in five other clusters associated with varied disease severity (Supp Fig 4). Thus, the unique association of this cluster with severe lung injury suggests a synergistic role of the E2F and MYB pathways during the severe phase of ARDS.

Lastly, the two distinct TSR clusters with the highest lung injury severity association have motif enrichment for STAT and NFkB (Modified Murray Score correlation, 0.518 and 0.417 respectively) (Fig 2b-c, top panel). Both clusters are enhancer-centric (71% and 62%) and enriched in neutrophil open chromatin (Supp Fig 3). Motif analysis demonstrated that the STAT and NFkB clusters are co-enriched, respectively, with BCL6 and RBPJ, the transcriptional effector for the NOTCH signaling pathway (Fig 3c, top panel).

We further validated the TF and TSR cluster association through orthogonal approaches. Cross-referencing the active cistrome with available TF ChIP-seq datasets reveals that TSR clusters are generally enriched for ChIP-seq signals associated with the predicted TF motif (Supp Fig 5c). Furthermore, we tested whether target genes of each TSR cluster overlap with gene signatures from a systematic experimental perturbation with chemical and small molecules intended for drug repurposing and discovery (Connectivity Map, CMap) (29). We expected to find therapeutic candidates that target the TFs predicted for each TSR cluster.

Indeed, CMap analysis revealed multiple cell-cycling inhibitors for the target genes from the E2F/MYB cluster, BCL inhibitor and JAK/STAT inhibitor for the STAT/BCL6 cluster, TGFβ receptor inhibitors for the ARE/SMAD/AP1 cluster, and cortisone for the GRE cluster (Supp Table 2). Overall, the unbiased discovery of TSR clusters based on the dynamic transcriptional activity, when coupled with motif analysis, revealed TF-regulated biological pathways during severe SARS-CoV-2 infection.

### Natural history of the transcriptional factor program in survivors of severe COVID

We next sought to delineate the temporal trajectory of the immune TF program during severe COVID-19 infection. Utilizing a subset of survivors (N = 9, Supp Table 1) with severe COVID-19 and prolonged hospitalization for COVID-19 associated ARDS with longitudinal sampling, we characterized the natural history of peripheral immune transcriptional programs. Correlation coefficients for TSR activity relative to hospital admission overlaid onto the cistrome UMAP displayed a non-random distribution of TSRs characterized by their temporal response patterns to COVID-19 recovery (Fig 4a). Most TSRs in clusters corresponding to T1ISRE/STAT, NFkB/RBPJ, STAT/BCL6, and X-Box/CRE/KLF/NFY displayed strong activation early in the hospitalized clinical course (Fig 4b-d). In contrast, the temporal correlation coefficients for TSRs in CEBP/PU.1, the promoter-centric YY1/CEBP/PU.1, and the monocytic ARE/AP1/SMAD are associated with late activation in the clinical course (Fig 4e). We observed consistent temporal patterns when the median TSR activity for each cluster was plotted across hospitalization time (Fig 4b-e). As independent corroboration of T1ISRE activity, the T1ISRE/STAT csRNA clustering paralleled the circulatory level of IP-10 (Fig 4f, Pearson’s correlation = 0.846, p-value < 1 x 10^4^), an interferon-induced cytokine associates with severe COVID-19 (5). STAT/BCL6 TF activity was also correlated with IL-6 plasma cytokine levels (Fig 4g, Pearson’s correlation = 0.641, p-value < 1 x 10^4^).

The inverse temporal profile of the neutrophil TSRs indicates a transition within neutrophil cistrome activity during recovery from COVID-19 ARDS. We performed gene ontology analysis on the target genes identified within temporal clusters. The target genes associated with the early neutrophil TF programs were enriched in interferon signaling, leukocyte activation, NFkB/TNFα signaling, autophagy, protein catabolic process, and membrane trafficking (Fig 4h, Supp Table 3). The target genes for the late neutrophil TF programs enriched in pathways associated with RNA splicing, translation, and protein and organelle localization (Fig 4h, Supp Table 3). TGFβ signaling was the top pathway in the monocytic clusters with ARE/AP-1/SMAD motif enrichment (Fig 3c). Notably, NFkB motifs were significantly depleted in both CEBP/PU.1 (Cluster 5) and YY1/CEBP/PU.1(Cluster 18) network, suggesting that the NFkB pathway was inactive during recovery (Supp Fig 4).

In summary, cistrome analysis across the clinical course identified a transition in TF network activity in the transcriptional program during the recovery of critically ill patients with COVID-19 ARDS.

### Distinct TF regulatory networks overlap genetic variants associated with COVID-19 clinical outcomes

We hypothesized that the regulatory elements we identified as associated with severe COVID-19 ARDS might overlap with genetic variants associated with COVID-19 outcome. To this end, we cross-referenced the active cistrome with disease-associated SNPs from the COVID-19 Host Genetic Initiative Consortium (30). The consortium performed a meta-analysis with a combined 49,562 COVID-19 cases identifying thousands of SNPs (p-value < 5 x 10^-8^) associated with COVID-19 clinical outcomes. We first identified the disease-associated SNPs within -300 to +100 bp of the TSS (Fig 5a-b). We tested for enrichment of disease-associated SNPs within active cistrome regulatory patterns using RELI (31), which assesses the significance of overlap between genetic variants and regulatory elements while considering the underlying genetic structure of the data. We focused on SNPs associated with hospitalization, with non-hospitalized COVID-19 cases as controls, mirroring the focus of our COVID-19 ARDS cistromic data. TSRs positively correlated with lung injury index (Modified Murray Score coefficient > 0.15) have significant enrichment for disease-associated SNPs associated with hospitalization (Padj 2.57 x 10^-13^) (Fig 5c). We next cross-referenced disease-associated SNPs with our TSR clusters identifying significant disease-associated SNP enrichment in distinct TSR clusters. The E2F/MYB, STAT/BCL6, and T1ISRE/STAT clusters are significantly enriched for COVID-19 hospitalization-associated SNPs when compared to non-hospitalized cases (Padj-7.62 x 10^-7^, 1.67 x 10^-9^, and 2.16 x 10^-13,^ respectively). Additionally, the three clusters with ARE motifs - MiT/NFkB/ARE, CEBP-AP1/ARE, and SMAD/AP-1/ARE (Padj- 6.2 x 10^-4^, 1.7 x 10^-3^, and 5.12 x 10 ^-11^, respectively) also have SNPs associated with COVID-19 hospitalization.

### Cistrome-disease relationship reveals dysregulated E2F/MYB, STAT/BCL6, and T1ISRE/STAT activity

Reflecting on the GWAS analysis, we hypothesized that the activity of TF-regulatory networks enriched for disease-associated genetic variants is likely correlated with disease outcome. Consistent with this notion, the two deceased patients in our cohort had persistently elevated STAT/BCL6 and E2F/MYB activities compared to survivors with a prolonged hospital course (Supp Fig 7). To extend these observations and test the predictive ability of our findings, we expanded our cistrome-based TF network analysis to measurements of gene expression (RNA-seq), under the assumption that target gene expression can quantify TF activity indirectly (Fig 6a). We focused on the E2F/MYB, STAT/BCL6, and T1ISRE/STAT TF regulatory networks because of their early activation in the hospital courses, consistent with most published COVID- 19 studies with transcriptomic data. Toward this goal, we used samples with matched cistrome and transcriptome data (n = 55) to identify E2F/MYB, STAT/BCL6, and T1ISRE/STAT target genes (n = 106, 176, and 58 respectively, see method). The average target gene expression for E2F/MYB, STAT/BCL6, and T1ISRE/STAT is highly correlated with the average TSR activity for those clusters (r = 0.92, 0.89, and 0.93 respectively) (Fig 6b, Supp Fig 8), enabling us to estimate the activity of these regulatory programs from blood leukocytes RNA-seq from individual patients.

Using this approach, we evaluated the correlation of E2F/MYB, STAT/BCL6, and T1ISRE/STAT network activity to disease severity in a large COVID-19 cohort (n = 100) with available peripheral leukocyte transcriptomics (32). This cohort used the number of hospital-free days at the 45th day of admission (HFD45) to delineate clinical severity – severe cases with prolonged hospitalization have fewer hospital-free days. Both E2F/MYB and STAT/BCL6 activity independently correlated with disease severity, with higher TF-network activity corresponding to lower number of hospital-free days (Fig 6c). T1ISRE/STAT activity did not show a linear relationship with disease severity; rather, patients at each extreme of T1ISRE activity trended toward poor disease prognosis (Fig 6c).

We then queried whether TF-network interactions track disease severity. The cohort was divided into “high” and “low” groups based on TF-network activity. The STAT^hi^E2F^hi^ group included significantly more fatal cases (HFD45 = 0, *Chi*-square, p-value = .004, two-tailed) with the lowest HFD45 (Fig 6d-e). When delineated by T1ISRE/STAT and E2F/MYB activity, the four groups have no statistically significant differences in fatality numbers (Chi-square, p-value = 0.47, two-tailed) or HFD45 (ANOVA, adj p-value = 0.34) (Fig 6f). T1ISRE/STAT and STAT/BCL6 did exhibit significant interaction. The T1ISRE^lo^/STAT^hi^ group (n=14) included the highest proportion of fatalities and lowest average HDF45 score (Fig 6g). This finding is consistent with the current literature on interferon dysregulation in COVID-19 (3, 8, 33), but it uniquely emphasizes that patients with combined high STAT/BCL6 and low T1ISRE/STAT activity are the most vulnerable.

### The imbalance of STAT/BCL6, E2F/MYB, and T1ISRE/STAT reflects the imbalance of neutrophil subsets

With emerging clinical correlates of immune subsets with COVID-19 severity, we investigated the relationship of the cistrome-signature with immune subpopulations. We cross-examined the enrichment of E2F/MYB, STAT/BCL6, and T1ISRE/STAT in COVID19 single-cell RNA-seq studies to see if these pathways converge on specific cell populations (8). Among all the leukocytes in the single-cell RNA-seq analysis, we found high enrichment of E2F/MYB, STAT/BCL6, and T1ISRE/STAT in neutrophils, comprising three subpopulations (Fig 6h, Supp Fig 9). The STAT/BCL6 signature is significantly enriched in all three neutrophil populations.

Notably, the subset consistent with immature neutrophils (LCN2^+^, CEBPE^+^) has the highest STAT/BCL6 enrichment, with nearly 50% of the transcripts per cell derived from target genes in the STAT/BCL6 network (Fig 6i). The immature neutrophil subset also has a high E2F/MYB and low T11SRE/STAT signature, resembling the expression signature associated with severe disease identified in both bulk RNA-seq cohorts (Fig 6i). The two other major populations are consistent with mature neutrophils, with each having approximately 20% of transcripts per cell derived from the STAT/BCL6 network. Notably, mature neutrophil 1 has the highest enrichment for target genes in the T1ISRE/STAT network, including numerous interferon-stimulated genes (ISGs), consistent with a subspecialized neutrophil population with antiviral activity (Fig 6i) (34). To track how these cell populations vary with the disease, we assessed cell counts in each patient stratified by disease severity, finding patients with COVID-19 have disproportionately higher neutrophil counts overall. However, patients with severe disease were more likely to have neutrophils from mature neutrophil 2 (*NEAT/S100A12*) and immature neutrophil subpopulations, implicating that STAT/BCL6 and E2F/MYB pathways likely converge to regulate the emergence of immature neutrophils in severe disease (Fig 6j).

## Discussion

We report the first study to profile initiating transcription in primary patient samples. One study has previously used nascent transcriptomics to identify active regulatory elements in patient samples (35), but our current study is the first longitudinal, active cistromic study of peripheral immune leukocytes in ARDS associated with COVID-19. Our analysis of the regulatory landscape catalogs the dynamic regulation of eRNAs and provides a TF-centric analysis and interpretation. Because the identity of the TF is revealed through the genomic DNA sequence, csRNA-seq coupled with motif analysis is, in essence, an unbiased functional assay for TF activity (36), and provides a novel dataset that is substantially different and complementary to traditional transcriptomics or other types of epigenetics profiling (e.g., ATAC-seq). Active cistromic analysis provides unique insights into the underlying TF networks and mechanisms in complex diseases when integrating with GWAS, bulk, and single-cell transcriptomics.

With this approach, we identified pathways and TFs implicated in severe COVID-19, including known therapeutic targets such as the glucocorticoid receptor, the interferon pathway (37), and targets currently in clinical trials including in the JAK/STAT pathway (38). Our analysis identified novel TFs in the immune response to severe COVID-19 and implicated their activity in neutrophils, including motifs for antioxidant response elements involved in oxidative homeostasis with NRF, small-MAF, and BACH. The genetic association of ARDS to NRF family *NFE2L2* in patients (21), and higher mortality due to bacterial pneumonia (39, 40) or more significant acute lung injury due to high tidal volume (41) in mice lacking *Nfe2l2*, supports the biological plausibility of our findings.

Identifying distinct co-TF regulatory networks is the principal finding of this study. With csRNA-seq and unsupervised clustering, we identified cis-regulatory elements and target genes with similar transcriptional initiation activity profiles clustering into distinct groups. Co-enrichment of TF motifs within a single cluster is therefore suggestive of cooperative regulation. This concept is classically exemplified by the interaction between signal-dependent and lineage-determining TFs, where cell-lineage pioneering TFs established chromatin accessibility for cell-type specific, signal-dependent regulation (22). Importantly, our dataset revealed networks with co-enrichment of multiple SDTFs, including 1) E2F and MYB, 2) MiT/TFE, ARE and NFkB, 3) NFkB and Notch, 4) STAT and BCL6, and 5) ARE/SMAD/AP1. The patient’s active immune cistrome thus provides evidence of unrecognized interactions between otherwise well-described pathways. MiT/TEF family of TF plays a crucial role in autophagy and lysosomal biogenesis for nutrient and energy homeostasis, adaptation to metabolic stress, and immune response (27, 42, 43). The convergence of ARE, NFkB, and MiT/TEF motifs in a single TF network suggests a biological significance in the interaction of redox, inflammation, and autophagy during COVID-19 ARDS. Similarly, the co-enrichment of ARE/SMAD/AP1 late in the hospital course for prolonged survivors suggests co-regulation of TFs in the TGFβ and redox pathways. Such has been shown in macrophages during wound healing, where the expansion of cistromic co-occupancy was noted for Nfe2l2, Smad3, AP1 family Fos1, and NFkB upon simultaneous exposure to tissue damage signals (28). This finding provides a collaborative model where TFs of different families converge in response to combinatorial biological signals in the cellular milieu.

The significant overlap of disease-associated SNPs in distinct TSR clusters suggests plausible functional importance of co-regulatory TF networks in COVID-19 outcome. Our cistrome dataset is uniquely complementary to the ongoing COVID-19 GWAS effort. The dataset directly identifies transcriptionally active genomic regions from immune cells from COVID-19 patients, with detailed annotation of disease severity and TF regulatory pattern associations. While a disease-associated SNP localizes disease risk to a genomic locus, enrichment of disease SNPs within multiple regulatory elements of a TF program implicates the biological significance of the TF pathway (31). We identified over-representations of COVID-19 associated SNPs at TSRs regulated by the T1ISRE/STAT, STAT/BCL6, and E2F/MYB networks. We further demonstrated that the activity of these TF regulatory networks parallels disease outcomes. Specifically, a combination of high STAT/BCL6 and E2F/MYB signals, even early in the hospital course, is associated with a poor prognosis. COVID-19 patients with low Type 1 interferon and high STAT/BCL6 activity also exhibited worse outcomes. Furthermore, these TF network signatures mapped to distinct neutrophil subsets. While our dataset cannot provide evidence of causal genetic variants, integrating the cistrome, GWAS, and transcriptomic analyses supports functional roles for co-regulatory TF networks.

Our work has several limitations. First, the study design maximizes temporal resolution, which limits patient numbers. A dedicated study with a larger cohort for csRNA-seq would be ideal for confirmation. Nonetheless, the validation analysis using a large external COVID-19 cohort confirmed the cistrome association with poor disease outcomes. It also demonstrates the feasibility of identifying target genes as a proxy for TF network activity. Secondly, we profiled the cistrome of all peripheral leukocytes, a heterogeneous cellular population with different proportions. The imbalance in cellular proportion influences clustering resolution, which is more sensitive to cells making up the majority of the heterogeneous population. Cell sorting prior to cistrome analysis would address this issue but presents technical and feasibility challenges requiring larger blood volumes from clinically unstable patients. To address cell-type identity, we cross-examined publicly available cistrome databases and successfully identified major inflammatory pathways in smaller subsets of circulatory immune cells, including the lymphocytic NFkB and the monocytic ARE/SMAD/AP1 programs. We also identified the combined STAT/BCL6 and E2F/MYB signature from immature neutrophils, which usually represent < 10- 20% of total leukocytes even in critical illnesses.

In summary, we used a novel unbiased technique to examine active genomic regulatory elements by profiling levels of initiating transcripts directly from peripheral leukocytes of critically ill COVID-19 patients. We identified and provided evidence of TF pathways and co-regulatory mechanisms implicated in severe COVID-19 ARDS. Many of these transcription factors are pharmacological targets for existing compounds. These pathways may also be critical players in infectious, non-COVID-19 ARDS, a heterogeneous clinical syndrome with high mortality (35- 40%) that currently depends on supportive care without targeted pharmacological therapy (44). We demonstrated the feasibility of using cistrome-derived TF networks to stratify patients by clinical outcome. Fast-turn around, TF activity profiling could be clinically applicable to stratify patients for TF-targeted therapy, such as anti-IL6 or JAK-STAT inhibitor for patients with high STAT/BCL6 activity. Equally important, one may avoid non-specific JAK-STAT inhibitors in severe patients with high STAT/BCL6 and low T1ISRE/STAT activity, where further impairment in the type 1 interferon pathway could be detrimental. Overall, our study demonstrates that unbiased active cistrome profiling offers an unprecedented TF-centric resolution in understanding human disease.

## Materials and Methods

### Study and Participants

Hospitalized patients diagnosed with COVID-19 at UCSD hospitals including Hillcrest and Jacobs Medical Centers as well as Rady Children’s Hospital were recruited for these studies from April to June 2020. After informed consent, blood was drawn on hospitalization days 1, 3, 5, 7, 9, 11 and discharge/death for analysis. Medical records were reviewed and patient demographics, laboratory values, and clinical characteristics were extracted using the Research Electronic Data Capture (REDCap) electronic data capture tool hosted at the University of California, San Diego.

### Characterizing Lung Injury with Modified Murray Score

The Murray Score was developed to characterize the level of lung injury in acute respiratory distress syndrome (15). This system assigned a score of 0-4 to the following 4 categories. 1) the extent of lung involvement on chest radiograph; 2) level of hypoxemia using PaO2 to FiO2 ratio; 3) the range of positive end expiratory pressure (PEEP); and 4) range of lung compliances based on tidal volume, peak inspiratory pressure, and PEEP. For this study, in order to stratify lung injury of patients prior to mechanical ventilation, after liberation from mechanical ventilation as well as requirement for advance therapy on mechanical ventilation, we added the mode of respiratory support. Patients on room air will be given 0 point; 1 point for 1-6 liter (L) of supplemental oxygen through nasal cannula; 2 points for non-rebreather mask at 10-15L of supplemental oxygen; 3 points for mechanical ventilation; and 4 points for mechanical ventilation with proning and paralysis. The modified Murray Score was tabulated by averaging the score from these five categories. A score of 0.1 to 2.5 was considered mild-moderate disease. Severe ARDS is > 2.5. Non-invasive positive pressure ventilations including bi-level and heated high flow nasal cannula were not included in the modified Murray score because at the time of recruitment, the safety of these modalities for exposing medical staff were not well understood, and their use was generally discouraged.

### Blood sample processing

Blood (3-10mL) was collected in a Sodium Heparin (BD Vacutainer, REF: 366480) or Potassium EDTA (BD Vacutainer, REF 367861) tubes. To prevent coagulation, the tubes were inverted 10 times prior to transport at room temperature. Blood was processed within 4 hours of collection and kept at room temperature throughout the protocol.

Whole blood from EDTA tubes and Heparin tubes was processed for isolation of plasma and whole white blood cells (WBC).

### Peripheral Leukocytes Isolation

To isolate WBCs, whole blood in EDTA tubes was centrifuged at 300xg for 20 minutes. Plasma was first removed, leaving a cell pellet containing WBCs and red blood cells (RBCs). RBCs were lysed by resuspending the cellular pellet in 1X RBC Lysis Buffer (ammonium chloride (8.02 g/L), sodium bicarbonate (0.84 g/L), and EDTA (0.37 g/L) in deionized water) and incubated for 10 minutes. The cell suspension was then centrifuged at 600xg for 5 minutes and the pellet was again resuspended in RBC lysis buffer for 5 minutes. The reaction was quenched with 3 times the volume of 1X HBSS (Gibco, REF 14175-095). After a sample was collected for a cell count, the isolated WBCs were pelleted at 600xg for 5 minutes. The WBC pellet was lysed in Trizol Reagent (Life Technologies, REF 15596018) with a target concentration of 5-10 million cells/mL and stored at -80C prior to RNA extraction.

### Plasma Isolation

Plasma for cytokine analysis was collected from Sodium Heparin tube. Plasma was removed from blood separated by Polymorphprep^TM^ per manufacturer’s instructions (Progen). Plasma was transferred to new microcentrifuge tubes and centrifuged at 3731xg for 5 minutes at room temperature to remove any cellular debris. Supernatant was transferred to new tubes and flash frozen in dry ice and 95% ethanol. Plasma was stored at - 80C for further analysis.

### RNA extraction and purification

Total RNA was extracted from WBCs using TRIzol(tm) reagent (Cat. No. 15596018, ThermoFisher Scientific) as per manufacturer’s instructions. Half of the total RNA was submitted for capped-small RNA-seq library generation. The remaining RNA was treated with TURBO(tm) DNase (AM1907, ThermoFisher Scientific) as per manufacturer’s instructions and used for bulk total RNA-sequencing.

### Capped small RNA-sequencing

csRNA-seq was performed as described in previously (11). Briefly, small RNAs of 20-65 nt were size selected from 0.3-1.0 microgram of total RNA by denaturing gel electrophoresis. A 10% input sample was taken aside, and the remainder enriched for 5’-capped RNAs with 3’-OH representing RNAPII initiated RNAs.

Monophosphorylated RNAs were selectively degraded by Terminator 5’-phosphate-dependent exonuclease (Lucigen). Subsequent 5’ dephosphorylation by CIP (NEB) followed by decapping with RppH (NEB) augments Cap-specific 5’ adapter ligation by T4 RNA ligase 1(NEB). The 3’ adapter was ligated using truncated T4 RNA ligase 2 (NEB) without prior 3’ repair to select against degraded RNA fragments. Following cDNA synthesis, libraries were amplified for 11-14 cycles and sequenced SE75 on the Illumina NextSeq 500 sequencer.

Sequencing reads were trimmed for 3’ adapter sequences using HOMER (“homerTools trim -3 AGATCGGAAGAGCACACGTCT -mis 2 -minMatchLength 4 -min 20”) and aligned to the human GRCh38/hg38 genome using STAR (45) with default parameters. Only reads with a single, unique alignment (MAPQ >=10) were considered in the downstream analysis.

Furthermore, reads with spliced or soft clipped alignments were discarded (the latter often removes erroneous snRNA alignments). Transcription Start Regions (TSRs), representing loci with significant transcription initiation activity (i.e. ‘peaks’ in csRNA-seq), were defined using HOMER’s findcsRNATSS.pl tool, which uses short input RNA-seq, traditional RNA-seq, and annotated gene locations to eliminate loci with csRNA-seq signal arising from non-initiating, high abundance RNAs that nonetheless are captured and sequenced by the method (full description is available in Duttke et al.(11). To lessen the impact of outlier samples across the data collected for this study, csRNA-seq samples were first pooled into a single META-experiment, and TSRs where then identified using findcsRNATSS.pl with a minimal TSR detection threshold of 1 read per 10 million mapped reads (“-ntagThreshold 1”), yielding 93,465 TSRs total. The resulting TSRs were then quantified in all samples by counting the 5’ ends of reads aligned at each TSR on the correct strand. The raw read count table was then normalized using DESeq2’s rlog variance stabilization method (46). The resulting normalized data was used for all downstream analysis. Normalized genome browser visualization tracks were generated using HOMER’s makeMultiWigHub.pl tool. TSR genomic DNA extraction, nucleotide frequency analysis relative to the primary TSS, general annotation, other basic analysis tasks were performed using HOMER’s annotatePeaks.pl function. Overlaps between TSRs and other genomic features (including peaks from published ATAC-seq studies, and annotation to the 5’ promoter of annotate GENCODE(v34) transcripts), was performed using HOMER’s mergePeaks tool.

### Total RNA sequencing

Libraries were prepared using Illumina’s TruSeq Stranded Total RNA Library Prep Gold according to manufacturer’s instructions. In brief, rRNA was depleted from total RNA (0.35 mg) by using subtractive hybridization. The RNA was then fragmented by metal-ion hydrolysis and subsequently converted to cDNA using SuperScript II. The cDNA was then end-repaired, adenylated, and ligated with Illumina sequencing adapters. Finally, the libraries were enriched by PCR amplification. All sequencing libraries were then quantified, pooled, and sequenced paired-end 150 base-pair (bp) on Illumina Novaseq at the Salk Next Generation Sequencing Core. Each library was sequenced on average 30 million reads. Sequencing reads were aligned to the human GRCh38/hg38 genome using STAR. STAR was also used to quantify read counts per gene using transcripts defined by GENCODE (version 34). RNA-seq read counts were then normalized using DESeq2’s rlog variance stabilization method (46) for all downstream analyses.

For total RNA-seq from Overmyer et al (47), sequencing reads were downloaded from GSE157103 and were processed in the same fashion (i.e. mapped with STAR, rlog normalized with DESeq2).

### Unsupervised machine learning for TSR cluster identification

We used hierarchical clustering and Uniform Manifold Approximation and Projection (UMAP) for unbiased clustering of TSRs based on csRNA-seq activity at the level of each TSR. First, patterns of csRNA-seq regulation were identified with unbiased hierarchical clustering using HOMER (“homerTools cluster”). The rlog normalized read counts across all TSRs were first row centered by the average read count for each TSR, and the data was subsequently hierarchically clustered using average linkage and the Pearson correlation coefficient between TSR profiles as the distance metric. Due to the size of the dataset, 10,000 random TSRs were first selected for hierarchical clustering. After completion, the remaining TSRs were assigned to their location in the hierarchical tree based on their nearest neighbor. The final clusters were defined as the maximum sized subclusters with an average correlation coefficient no greater than 0.30 and a minimum of 500 TSRs (to exclude small, highly variable clusters), yielding a total of 26. To visualize the data, we performed UMAP independently using R packages uwot, leiden, igraph, and FNN, with the following setting: n_component = 2, n_neighbors = 16, a = 2.5, b = 0.575, and metric = ‘correlation’. We used ggplot to visualize the UMAP projection and to overlay information including cluster IDs, relative csRNA-seq activity, chromatin accessibility, Modified Murray Score correlation coefficient, and Hospital Time correlation coefficient.

### Motif enrichment analysis

To identify TF motifs in TSRs that are associated with clinical scores and other quantitative phenotypes, we applied MEIRLOP, a tool that uses logistic regression to model the presence of motifs in a set of scored DNA sequences while accounting for simple nucleotide composition bias, such as that introduced by CpG Islands (20). TSRs were first scored by how well their activity profile across samples correlated (Pearson) with the Modified Murray Score, such that TSRs with relatively high activity in samples from very sick patients (i.e. high Modified Murray Score) yield high correlation coefficient values, while TSRs with relatively high activity in healthy patients yield low values. Sequences from -150 to +50 bp relative to the primary (mode) TSS within each TSR were then extracted, and these sequences and their associated correlation scores were then analyzed using MEIRLOP. 438 TF motifs in HOMER’s known motif database were evaluated and the motifs yielding the most extreme enrichment coefficients with significant p-values were reported.

To identify motifs associated with discrete TSR clusters, we used HOMER (22) to scan for DNA motifs in each TSR (-150,+50) using HOMER’s known motif database, assigning the presence of a motif to each TSR if the motif was detected at least once. Motif enrichment for each cluster was calculated by comparing motif occupancy rates in each cluster versus all other clusters to calculate the log2 enrichment and significance using the Fisher Exact test. The top enriched motif(s) were then used to label the clusters, accounting for highly similar motifs from large families.

### Integration of csRNA-seq data with hematopoietic NGS epigenome profiling

Previously published bulk epigenomics data (ATAC-seq/ChIP-seq) from isolated hematopoietic cell types was used to assess the potential cell-type specificity of TSRs identified in our whole leukocyte csRNA-seq profiling experiments. ATAC-seq data from GSE118189 (16) 26 different peripheral blood cell types was further supplemented with ATAC-seq from primary neutrophils from GSE150018 (17) to analyze open chromatin. H3K27ac ChIP-seq data for 21 different peripheral blood cell types were downloaded from the Blueprint Epigenome project (https://www.blueprint-epigenome.eu/) (18) to assess regions with active chromatin modifications. TF ChIP-seq data for RELA(NFkB) in monocyte derived macrophages (MDM) (GSE100381 (24)), NFKB1 in CD4+ T cells (GSE116695 (25)), STAT3 in MDM (GSE120943 (48)), GR in MDM (GSE109438 (49)), and IRF1 in MDM (GSE43036 (50)) were used to confirm the binding of TFs to predicted sites based on DNA motif analysis of TSR sequence. For ATAC-seq and TF ChIP-seq experiments, sequencing reads were downloaded from NCBI SRA, trimmed for adapter sequences, and aligned to the hg38 genome using STAR with default parameters. Replicate experiments were pooled by concatenating alignment files. Uniquely aligned reads (MAPQ>10) were then analyzed using HOMER to find peaks using “-style atac” and “-style factor” for ATAC-seq and TF ChIP-seq experiments, respectively. TF ChIP-seq peaks were found using their respective negative control input sequencing experiments, while ATAC-seq peaks were found using the pooled input from all ChIP-seq experiments as a control. HOMER was used to create genome browser tracks, quantify reads in the vicinity of TSRs to quantify enrichment, and create histograms of read distributions relative to the primary TSS (e.g. Supp. Fig. 1c). H3K27ac ChIP-seq data from the Blueprint Epigenome project was downloaded as bigWig files, converted to bedGraph files using UCSC’s bigWigToBedGraph utility, and quantified at TSRs using HOMER’s annotatePeaks.pl program using the ‘-bedGraph’ option.

To score TSRs by their cell type-specific ATAC-seq enrichment (i.e. Fig. 2c), ATAC-seq reads (normalized to 10^7^ total mapped reads) for each cell type were quantified in the vicinity of all TSR (+/-200bp from the primary TSS or each TSR). For each TSR, the enrichment for each cell type was defined as the log2 ratio of reads from that cell type divided by the average normalized read count for all cell types. The same approach was used to score ChIP-seq specific enrichment (i.e. Supp. Fig. 3, 5c) by quantifying each ChIP-seq experiment across TSRs (+/-200 bp for TF ChIP-seq, +/- 500 bp for H3K27ac ChIP-seq). Aggregate cluster cell-type enrichments were reported by calculating the average TSR cell-type specific enrichment for each of the TSRs in the cluster/TF-network (i.e. Supp. Fig. 3, 5c). Cell type enrichment patterns were further hierarchically clustered using Cluster 3.0 (51) (Pearson Correlation, average linkage) and visualized using Java TreeView (52).

### Overlap of disease-associated SNPs from COVID-19 GWAS with TSRs

GWAS meta- analysis results from the COVID-19 Human Genetics Initiative (30) corresponding the A2, B1, B2, and C2 comparisons (Version 6, hg38 version) were downloaded from the consortium website (https://www.covid19hg.org/results/r6/). Significant disease-associated SNPs were defined using a p-value threshold of 5x10^-8^ as recommended in the original study (30). To visualize disease-associated SNPs overlapping TSRs in the TSR UMAP, we identified the SNP with the most significant p-value that overlapped each TSR within -300 to +100 relative to the primary TSS.

Regulatory Element Locus Intersection (RELI) (31) was used to assess the significance of overlap between lists of TSRs (e.g. TF clusters, TSRs with modified Murray Scores > 0.15, etc.) and significant GWAS SNPs. First, lists TSRs were first mapped from the hg38 to the hg19 version of the human genome using UCSC’s liftOver tool. Next, lists of significant disease-associated SNPs were expanded to include additional SNPs in LD (r^2^ > 0.8) using SNIPA (EU ancestry) (53). These data were then used to run RELI using default parameters, looking for

SNPs that overlapped TSRs from -300 to +100 relative to the primary TSS, reporting the corrected p-value and overlapping SNPs (Supp. Table 7).

### Target genes selection and pathway analysis

Traditionally target genes are identified as the nearby genes to regulatory elements (54). Because csRNA-seq simultaneously profiles transcriptional initiation of protein-coding genes and cis-regulatory elements, the initiation activities of the target genes should correlate with the cis-regulatory elements. We defined target genes if they have promoter TSR that are in the same TF-network cluster. These target genes were submitted for Gene Ontology and Pathway Analysis using Metascape (55).

### Calculating TF-network activity from csRNA-seq

The aggregate TSR csRNA-seq signal represents the activity of the TF-network program. For each TSR cluster, the median csRNA-seq normalized count (log2) for all TSRs in a given cluster was calculated to represent TF-network activity.

### Target gene selection to calculate TF-network activity from transcriptomes

To compute TF-network activity from transcriptomic data, we first need to identify target genes whose steady state RNA levels reflect regulation at the transcription level. The target genes RNA level from total RNA-seq should match the transcriptional initiation activity from csRNA-seq. We used matching samples to process both total RNA-seq and csRNA-seq (n = 55) to identify target genes within the E2F/MYB, STAT/BCL6, and T1ISRE/STAT network. We computed a Pearson correlation coefficient for each target gene’s RNA level to the activity pattern of the cistromic TSR cluster across the 55 samples. Target gene was then selected by the following criteria: 1) the gene has a TSR resides in the annotated promoter and clusters into the E2F/MYB, STAT/BCL6, or T1ISRE/STAT network; 2) the gene is unique to one TF-network; 3) the gene is expressed in the transcriptomic analysis, defined by having a median normalized read count greater than 4 FKPM across all samples; 4) the target genes RNA level is positively correlated with the activity of the TSR cluster (correlation coefficient > 0.2). The last criteria remove genes whose steady-state RNA level can be influenced by post-transcriptional mechanisms. We identified 106, 176, and 58 target genes for E2F/MYB, STAT/BCL6, and T1ISRE/STAT respectively. For validation, the TF-network activity determined by TSRs (csRNA-seq) and target genes (total RNA-seq) for the three TF-networks were analyzed in a 3x3 correlation matrix with Pearson’s correlation and two-tail test for statistical significance (Prism 9).

### Integrated analysis with single cell RNA-seq data

Single-cell RNA-seq data from Combes et al (8) was analyzed using the SCANPY toolkit (56). Cells with greater than 20% mitochondrial or 50% ribosomal RNA reads were excluded, as were cells with fewer than 100 genes detected. In some cases, previous annotation labeling cells as possible multiplets was available and used to filter out non-singlets. Gene set signatures were tabulated per cell from raw read counts. For cell type identification, read counts per cell were normalized and log-transformed before applying the regress_out function to the total counts. The counts were then scaled to unit variance and zero mean. The cells were run through PCA, neighbor graph generation, and UMAP with default parameters. Cell clusters were identified using Leiden clustering with a resolution parameter of 0.25. Marker genes from these clusters were used to identify five neutrophil clusters.

### Cytokine measurements

The Human Anti-Virus Response Panel (13-plex; BioLegend, San Diego, CA) was used to quantitate human plasma cytokines (IL-1β, IL-6, IL-8, IL-10, TNFα, and IP-10). Plasma samples were stored at ^-^80°C until use. For the cytokine assay, plasma was freshly thawed at room temperature, centrifuged at 1,000 x g for 5 min, and run at a 2-fold dilution in Assay Buffer per the manufacturer’s instructions. Samples were acquired on a Canto II flow cytometer (BD) using a high throughput sampler. Samples were run in duplicate unless plasma volume was inadequate, and standards were run on all plates. Cell signaling technology (CST) was run prior to all flow cytometry runs to ensure low detector CVs and set laser delay.

### LEGENDplex Data Analysis Software (BioLegend) was used for analysis

#### Drug repurposing & connectivity mapping

CMap (https://clue.io/cmap) provides expression similarity scores for a specific expression profile with other drug-induced transcriptional profiles, including consensus transcriptional signatures of 2,837 drugs grouped into 83 drug classes (29). The connectivity score from CMap is calculated based on the observed enrichment scores in the queried gene lists relative to transcriptional signatures in the L1000 reference database. The score incorporates a nominal p-value calculated based on the comparison between the query and reference signatures relative to a null distribution of random queries, using the Kolmogorov-Smirnov enrichment statistic, which is then corrected for multiple testing using the false discovery rate method (29, 57–59).

For drug repurposing, the connectivity map scores were computed based on the target genes for each TF-network cluster. We hypothesized that the gene expression pattern resulting from the perturbation by a therapeutic compound should negatively correlate with the COVID-19 transcriptional signature as previously shown (58, 59). Therefore, we selected those compounds that had significant negative connectivity map scores (i.e. compounds with the connectivity scores *< −*90, predicted to reverse our input signature (29)). For each cluster, we grouped predicted drugs into: (1) individual drug lists with connectivity scores (cs) <-90; (2) Drugs for each cluster based on the predicted cluster targets with cs <-90; and (3) Pharmacological drug classes (i.e. JAK inhibitors) to determine which broader classes of drugs may be predicted by each cluster to reverse the COVID-19 transcriptional signature (cs <- 90).

#### Statistical Analysis

The appropriate statistical analysis is performed and detailed in the corresponding sections above.

#### Study Approval

The study was approved by the Institutional Review Board at the University of California, San Diego (UCSD IRB#190699). Written informed consent was received prior to patient’s participation in the study.

## Supporting information

Lam et al suppl table 7

Lam et al suppl information

## Author Contributions

C.W.B., M.T.Y.L., and N.G.C. conceived the study. J.F. and N.G.C. oversee the logistics of the trial. M.F.O., M.T.Y.L., and N.G.C. identified, enrolled, and consented patients. M.F.O., M.T.Y.L., M.R., M.H., M.A, K.P. collected patients samples. E.H., C.N., S.T., R.K., M.T.Y.L., S.D, and N.G.C.. processed patients samples. M.F.O., M.T.Y.L., A.P., L.C.A., K.P., and N.G.C. gathered and interpreted the clinical data. S.H.D. generated csRNA-seq libraries. H.L. generated total RNA-seq libraries. M.W.C. processed the sequencing data. C.W.B., M.W.C., M.T.Y.L., S.H.D. analyzed and interpreted the data from csRNA-seq and total RNA-seq. C.W.B. integrated csRNA-seq with reference epigenomes and TF-ChIP-seq. M.W.C performed the single cell RNA-seq analysis. S.I.R., J.M.D., R.K.A., and S.C performed the cytokine analysis. A.S.W. and N.G.C. performed CMap analysis and interpretation. A.M. and S.P. provided critical feedbacks on the study design and the manuscript. N.G.C., M.T.Y.L., and C.W.B. interpreted data and wrote the manuscript with input from all authors.

## Acknowledgements

We are grateful to all the patients and their family for participation in the study. We are also thankful for the nurses in the UC San Diego Hilcrest Medical Center ICU and the step-down unit for their assistance. We thank Dr. Christopher K. Glass for his advice. This work was supported by the NGS Core Facility of the Salk Institute with funding from NIH-NCI CCSG: P30 014195, the Chapman Foundation and the Helmsley Charitable Trust. M.T.Y.L. is supported by the Academic Sleep Pulmonary Integrated Research/Clinical Fellowship through the American Thoracic Society and by the NIH (5T32HL134632-04). M.F.O. is supported by NIH/National Institute of General Medical Science (T32GM121318). R.K.A. is currently funded by NIH K99 AI145762. S.H.D. is supported by NIH (K99GM135515). C.W.B. is currently funded by NIH (U19A135972 and GM134366).This study was approved by the University of California, San Diego Institutional Review Board IRB-190699.

