## Supplementary material for "Profiling Transcription Initiation in Peripheral Leukocytes Reveals Severity-Associated Cis-Regulatory Elements in Critical COVID-19": Lam et al suppl information

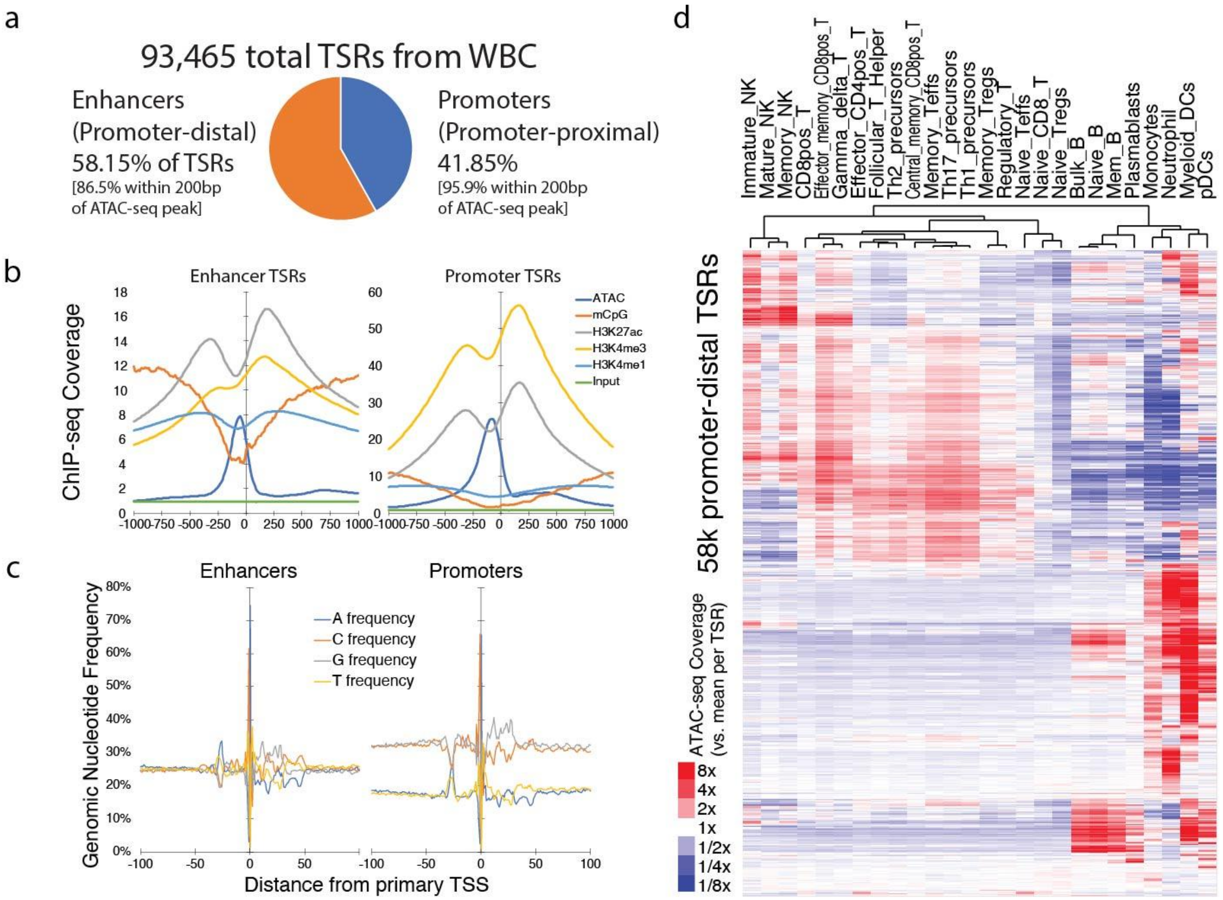


Supp Figure 1. The characteristics of Transcription Start site Regions (TSRs) identified by csRNA-seq from the peripheral leukocytes of COVID19 patients. a) 58.15% of the 93,465 transcription start site regions (TSRs) reside in promoter-distal regions. b) Epigenetic characteristics of TSRs located in promoter-distal regions are consistent with active enhancer signatures (Left) consisting of high acetylation of H3K27, and high H3K4me1 with lower H3K4me3 compared to TSRs in promoter regions (Right). Epigenetics data are aggregated across cell types from the Blueprint Epigenomics project [1]. c) Distinct distribution of genomic nucleotide frequencies flanking the TSRs in the enhancers (Left) and promoters (Right). The increased T and A frequency between -20 to -30 bp from the transcription start site (TSS) corresponds to the TATA box. E) The cell-type specific chromatin accessibility of flow-sorted peripheral immune cells at the promoter-distal TSRs [2,3]. Red depicts a higher ATAC-seq signal than the mean signal at each TSR; blue indicates ATAC-seq signal depletion.

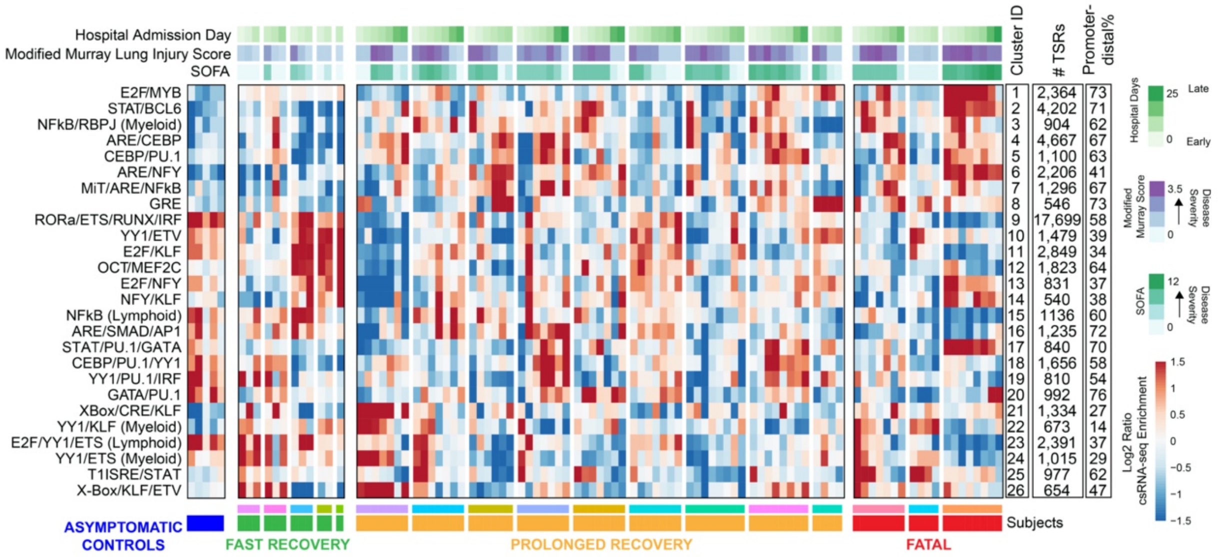


Supp Figure 2. Unsupervised clustering reveals discrete clusters of transcription start site regions (TSRs). Hierarchical clustering from 97 active cistromes revealed 26 discrete clusters with distinct activity patterns. Each row in the heatmap represents the relative average activity of a TSR cluster determined by csRNA-seq. Red indicates high enrichment relative to the mean of each row; Blue, low enrichment. Each column represents a single sample. The clusters are labeled with the highest TF-motif enrichments. Peripheral leukocytes were collected from asymptomatic controls (blue, n = 5), and COVID-19 patients with fast recovery (n = 5), prolonged recovery (n = 9), and fatal outcome (n = 3). Peripheral leukocytes were profiled at different time points in the hospital course with varying clinical status as indicated by the lung injury index (Modified Murray Score) and SOFA score.

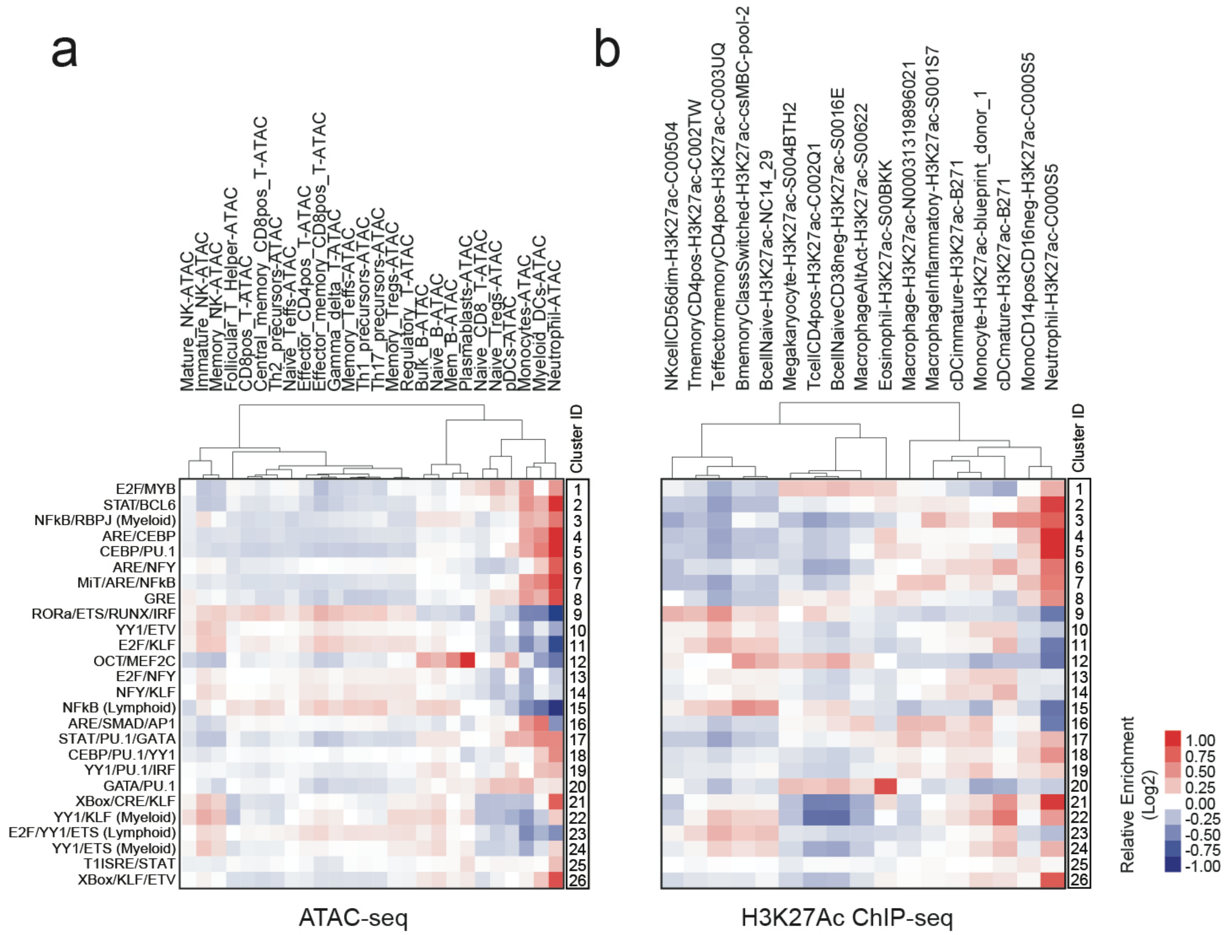


Supp Figure 3. Cell-type specific enrichment of chromatin accessibility and activated histone marks in TSR clusters from peripheral leukocytes from patients with active SARS-CoV-2 infection. Average signal from each cluster for a) ATAC-seq and b) H3K27ac ChIP-seq from flow-sorted peripheral immune cells [1-3] are depicted in the heatmap, with red showing higher signal enrichment and blue for lower enrichment.

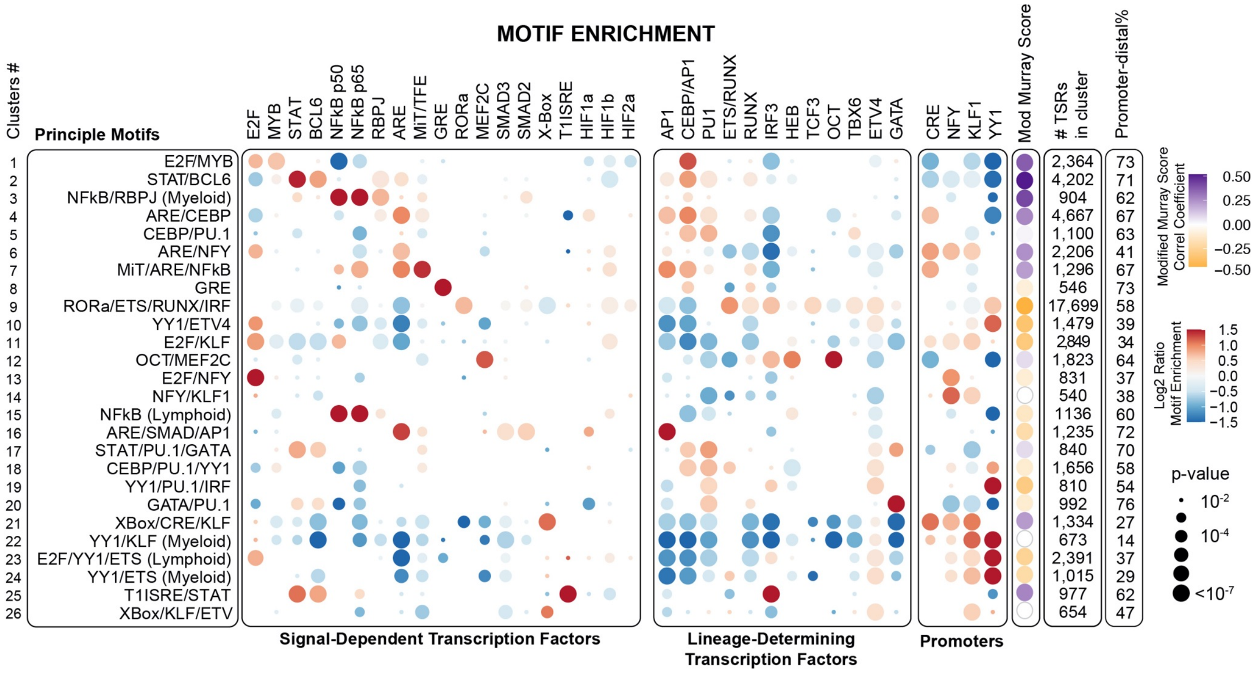


Supp Figure 4. Agnostic clustering of the active peripheral leukocytes cistrome identified TSR clusters with multiple TF motif co-enrichment suggestive of co-regulatory mechanisms. We searched for TF motifs from -150 to +50 bp from the primary transcription start site in each TSR. Enrichment analysis shows the frequency of TF motifs in each of the 26 discrete TSR clusters relative to all 93,465 active TSRs. Observed frequencies were expressed as Log2 ratio over the background. Red indicates a higher than expected TF motif frequency; blue, lower than expected. A larger bubble indicates lower Fisher's Exact test p-value (two tailed). The Modified Murray Score correlation coefficient depicts the relationship of the cluster activity with lung injury. Purple indicates positive disease correlation; orange, negative correlation.

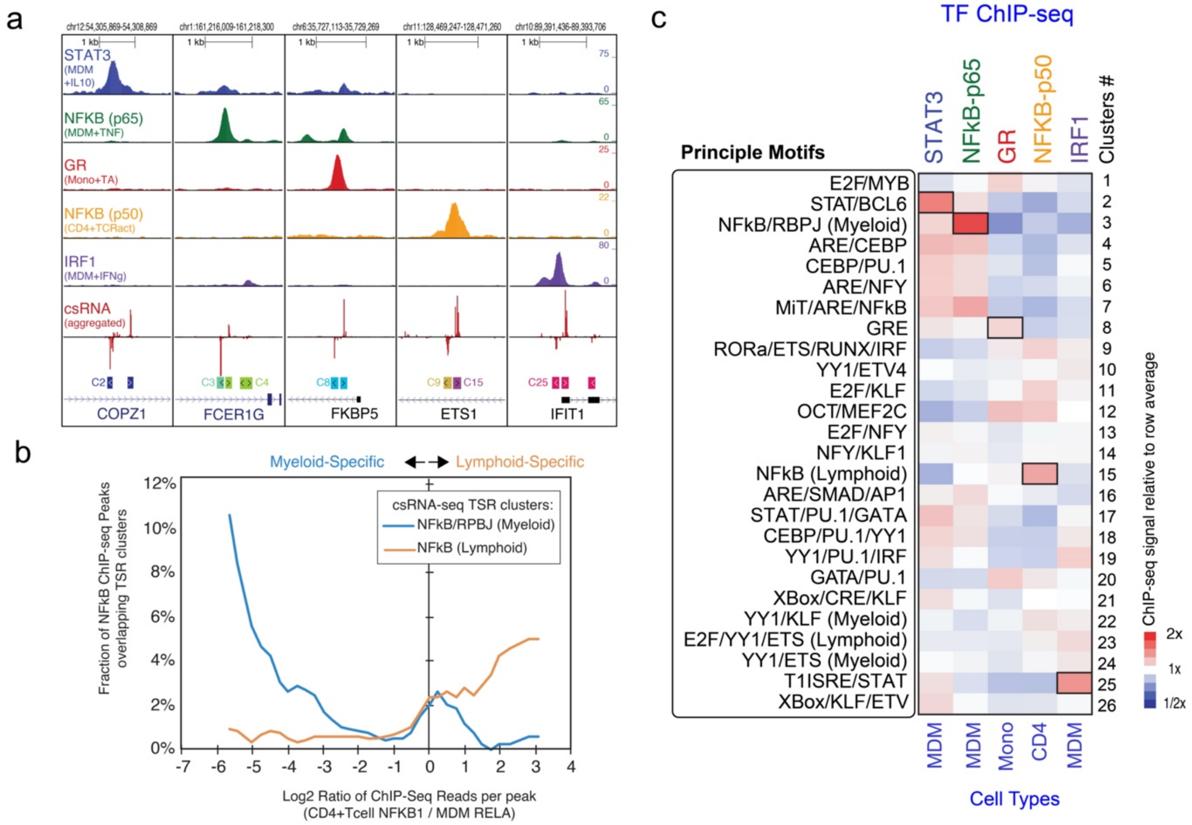


Supp Fig 5. Motif enrichment in TSR clusters overlaps with corresponding TF ChIP-seq signals. a) Five genomic loci with csRNA signal (lower track) overlap specifically with ChIP-seq for STAT3 [4] in the *COPZ1* locus, p65 [5] in the *FCER1G* locus, Glucocorticoid Receptor (GR) [6] in the *FKBP5* locus, p50 [7] in the *ETS1* locus, and IRF1[8] in the *IFIT1* locus. Cluster 2 (C2) was enriched for STAT and BCL6 motif; C3 for NFkB/RBPJ motifs; C8 for GRE motifs; C15 for NFkB motif in lymphoid lineage; C25 for T1ISRE/STAT. b) Cell-type specific NFkB chromatin localization in myeloid- and lymphoid-associated NFkB TSR clusters. Higher proportion of TSRs from lymphoid-specific NFkB cluster overlap with ChIP-seq peaks that have a higher signal for NFkB ChIP-seq in activated CD4 T cells than in activated macrophages (Log2 ratio > 0, orange curve). Reciprocally, NFkB ChIP-seq peaks with greater ChIP-seq signal in macrophages than in CD4 T cells are more likely to overlap TSRs from the NFkB/RBPJ (Myeloid) cluster (Log2 Ratio < 0, Blue curve). c) Global relative enrichment of TF ChIP-seq signal at TSR clusters. Red depicts high enrichment; low indicated depletion, boxes indicate cluster-TF-ChIP associations predicted by motif analysis. MDM = monocyte-derived macrophages. Mono = Monocytes. TNF = Tumor Necrosis Factor. TCRact = T-cell receptor via CD3 CD28 activation. IFNg = interferon gamma. TA = Triamcinolone acetonide.

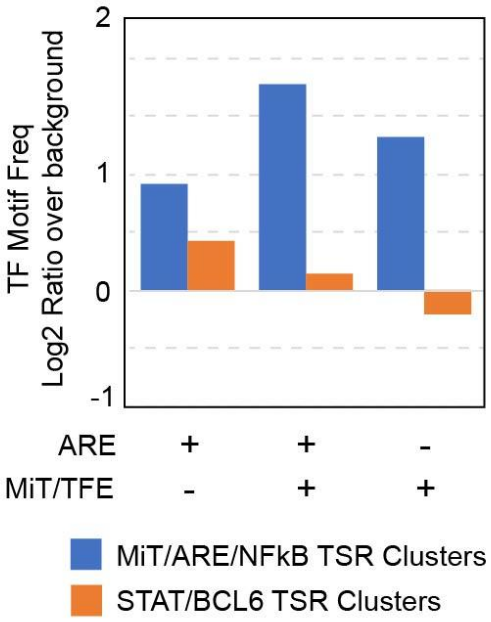


Supp Figure 6. Enriched co-occurrence of MiT and Antioxidant Responsive Element (ARE) motifs in the same TSRs in the MIT/ARE/NFkB cluster compared to background (*Chi*-square < 1 x $10^{-5}$, two tailed). As a control, the co-occurrence of MiT and ARE motifs is not enriched in the STAT/BCL6 TSR clusters. Log2 ratio is greater than 0 if the TF frequency is higher than background.

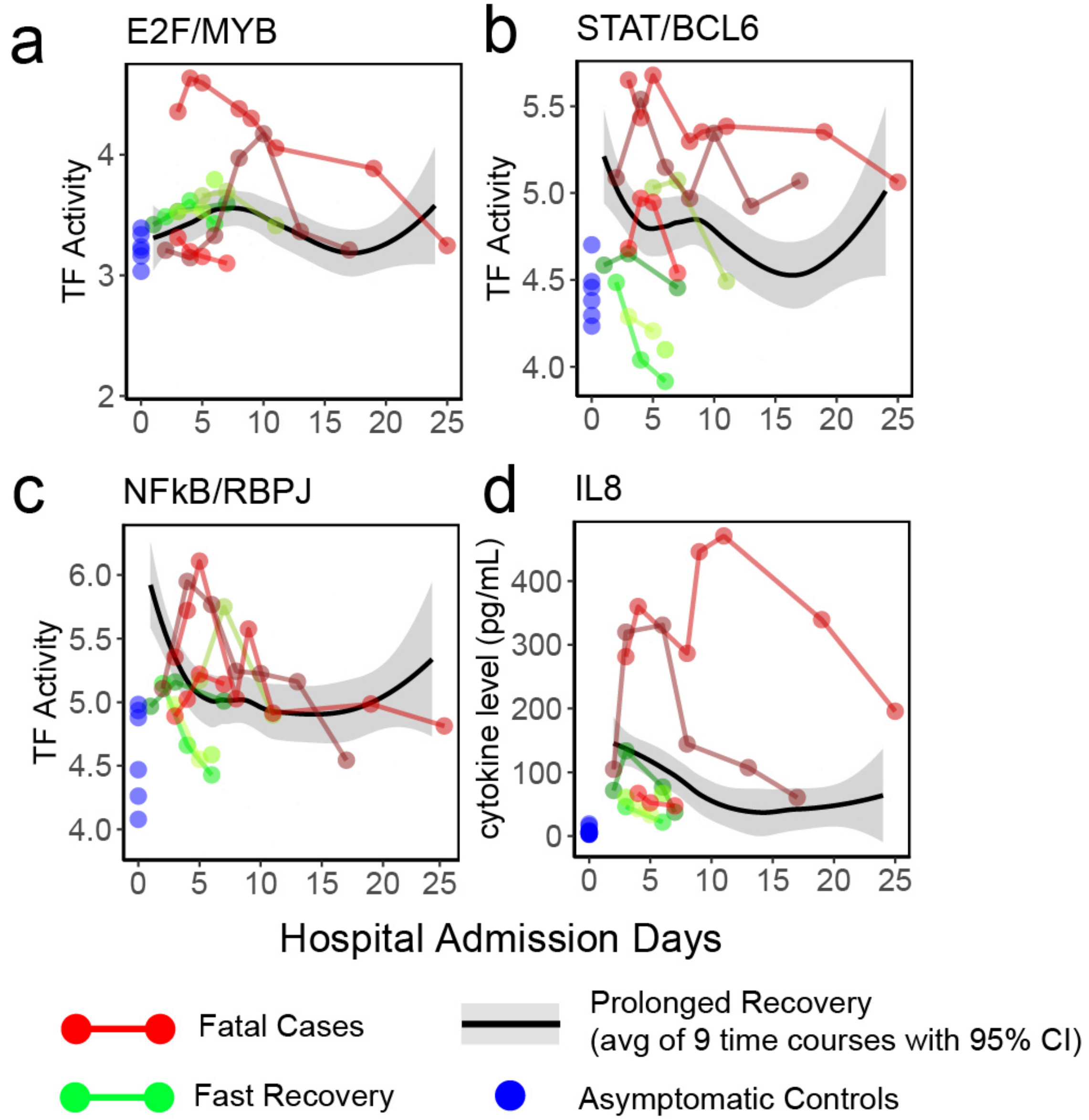


Supp Fig 7: Persistently elevated E2F/MYB and STAT/BCL6 activity in fatal COVID19 cases. Cistrome activity of A) E2F/MYB, B) STAT/BCL6, and C) NFkB/RBPJ programs with respect to the hospital course for individual patients with fatal outcomes (red) or fast recovery (green). The time course for 9 critically ill COVID19 patients with prolonged recovery is plotted as a smooth conditional mean with 95% confidence interval (gray shaded regions). D) Serum IL8 cytokine levels. The timing for cistrome and cytokine samples collections were within 24 hours.

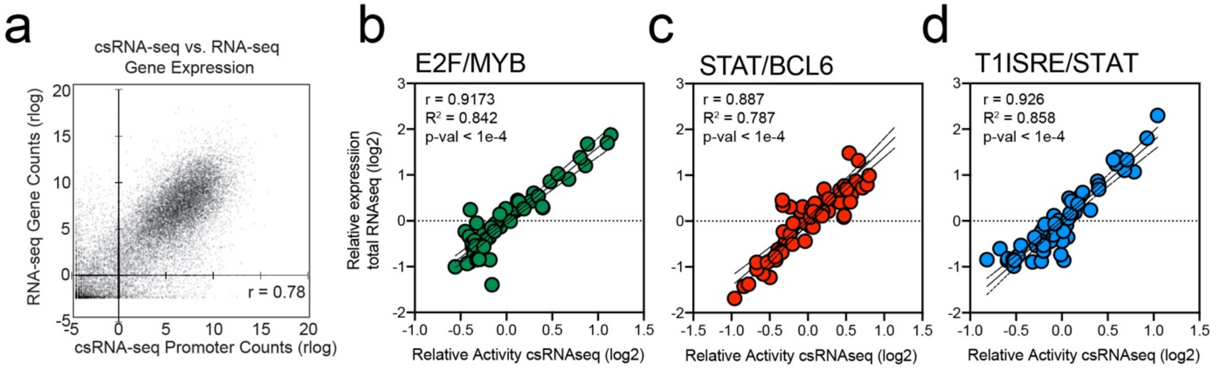


Supp Fig 8. Cistromic activity of E2F/MYB, STAT/BCL6, and T1ISRE/STAT represented by expression of TF target genes. a) Genome wide expression measured by csRNA-seq (x-axis) and total RNA-seq (y-axis) showed high correlation. b-d. Correlation analysis of b) E2F/MYB, c) STAT/BCL6, d) T1ISRE/STAT activities from 55 matched-samples with csRNA-seq and total-RNAseq profiling. Each point represents an average signal of TSRs (x-axis) and target gene expression (y-axis) in each TF program from matched samples. Statistical significance was determined by Pearson's correlation. r = Pearson's correlation coefficient. $R^2$ = coefficient of determination.

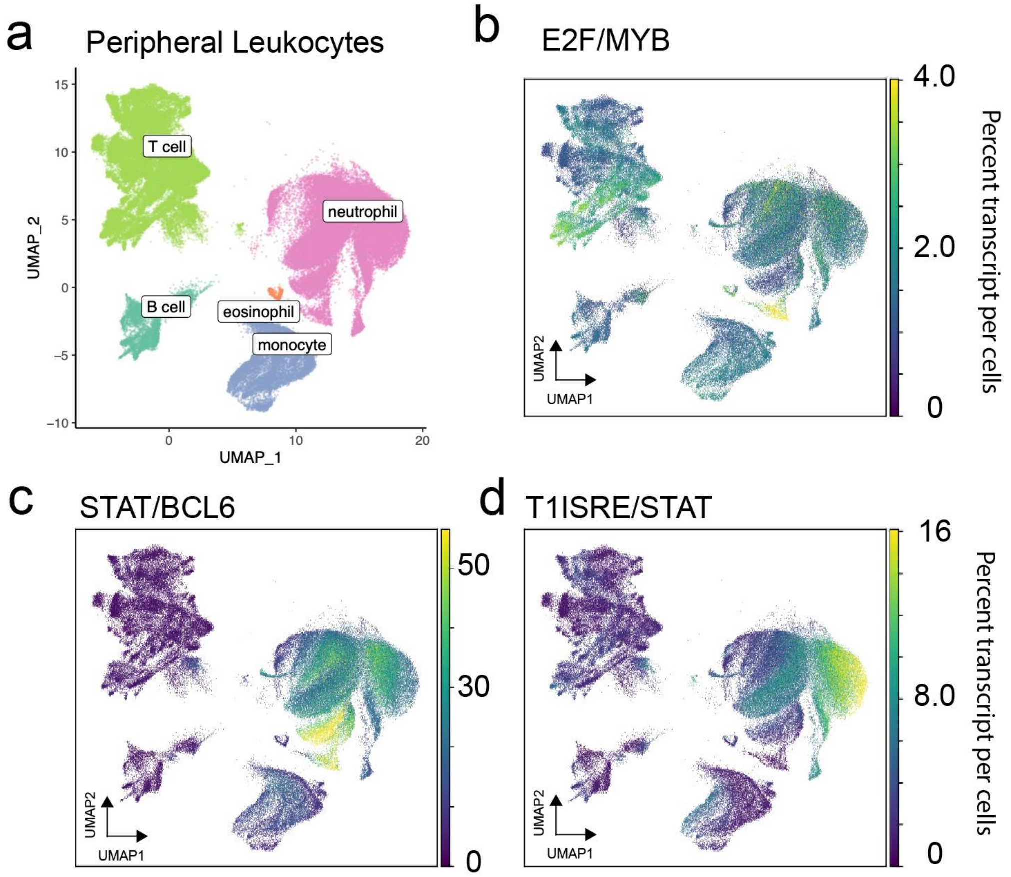


Supp Fig 9. E2F/MYB, STAT/BCL6, and T1ISRE/STAT activities converge in peripheral neutrophils. TF network activities based on the expression of target genes within the E2F/MYB, STAT/BCL6, and the T1ISRE/STAT networks were interrogated in a single cell RNA-seq dataset from peripheral leukocytes of a COVID-19 cohort [9]. a) Overall landscape of peripheral leukocytes clustered into major immune classes. b-d). The percent of transcripts per cell derived from b) E2F/MYB, c) STAT/BCL6, and d) T1ISRE/STAT TF network programs
